# Design and Evolution of an Orthogonal HaloTag for Multiplexed Labeling in Cells

**DOI:** 10.64898/2026.05.14.725131

**Authors:** Benjamin J. Goldberg, Bryan J. Lampkin, Alexander T. Stead, Siena Stanten, Patrick Rabe, Joshua A. Kritzer

## Abstract

The self-labeling protein HaloTag is used to install a wide variety of functional small molecules in cells and living organisms with exquisite specificity with respect to cell type and subcellular localization. HaloTag is a core part of many biotechnology-based tools for sensing, tracking, and manipulating biological systems with a high degree of spatial and temporal control. Due to the limitations of fluorescent proteins and other self-labeling proteins, most of these tools have historically been restricted to a single channel. In this work, we used structure-guided rational design and directed evolution to produce an orthogonal HaloTag protein called OrthoTag which reacts selectively with a modified chloroalkane substrate. OrthoTag retains many of HaloTag’s superior properties, and reaction rate measurements with purified proteins show OrthoTag and its substrate have 109-fold mutual orthogonality to HaloTag with respect to substrate concentration. We demonstrate the application of OrthoTag for multiplexed labeling experiments in mammalian cells with minimal optimization. Going forward, OrthoTag should be straightforward to incorporate into HaloTag-based systems to allow simultaneous measurement or manipulation of two biological targets or processes. The availability of multiple high-performance self-labeling proteins will enable the continued development of new multiplexed biotechnology methods.

## Introduction

A self-labeling protein (SLP) is an enzyme that catalyzes covalent bond formation between itself and a small molecule substrate.^1–4^ SLPs are most commonly used to attach synthetic probes to a fusion protein inside cells to mark, monitor, or modulate the fusion protein. When used with synthetic fluorescent dyes, SLPs overcome many of the limitations of fluorescent proteins for imaging in living systems due to the superior photophysical properties, stability, and tunability of synthetic dyes.^5,6^ Beyond fluorescent dyes, many different functional compounds have been attached to SLPs in live cells including drugs, reactive metabolites, metal catalysts, nucleic acids, and affinity handles.^7–19^ SLPs have enabled large advances in photoactivated proximity labeling, molecular recorders, and artificial metalloenzymes, and they are core parts of precision tools for studying cellular redox responses, complex neuropharmacology, and cytosolic penetration of large-molecule therapeutics.^7–19^ For all these diverse applications, HaloTag has consistently been used as the SLP of choice due to its superb reaction kinetics, its biorthogonality in eukaryotic cells, and the high permeability and ease-of-use of its chloroalkane substrate.^1,7–29^

To date, most HaloTag-dependent assays have not been multiplexed due to a lack of a suitable orthogonal SLP. Orthogonal SLPs to HaloTag, including SNAP-tag, CLIP-tag, and TMP-tag, have historically suffered from slower reaction kinetics, reduced substrate permeability, poor substrate washout, and unfavorable interactions with endogenous eukaryotic proteins.^2–4,22,29,30^ The recent development of SNAP-tag2 addresses some of these issues, though its optimized substrates still possess poorer predicted pharmacological properties compared to HaloTag’s substrate (Fig. S1).^31,32^ In the few examples where HaloTag was multiplexed, it was most commonly paired with conventional fluorescent proteins or luminescence reporters, which are less powerful for imaging and incompatible with many applications.^20,25^

Recently, we and others have successfully engineered HaloTag using structure-guided rational design, computational design, and/or yeast display.^15,16,20,25,33–35^ These efforts produced variants of the most widely-used SLP, HaloTag7,^28^ which have altered fluorescence lifetimes, improved reaction rates with specific dye-conjugated substrates, improved stability, and improved turn-on fluorescence with fluorogenic substrates.^31,33,34,36^ However, these variants all use the same linear chloroalkane ligand, and they require specific dye-protein interactions for some degree of orthogonal readout. In this work, we develop an orthogonal HaloTag7 variant (OrthoTag) and substrate, suitable for the simultaneous covalent labeling of two different proteins in live cells. By developing an orthogonal SLP that nonetheless has a large degree of homology with HaloTag7, we anticipated that OrthoTag would be readily implemented into existing HaloTag-based technologies, multiplexing them with minimal additional optimization.

## Results

### Bump-and-Hole Engineering of HaloTag

We reasoned that the “bump-and-hole” method could be used to generate an orthogonal HaloTag enzyme/substrate pair.^37–39^ Based on a crystal structure of HaloTag7 bound to a tetramethylrhodamine-conjugated substrate,^26^ we identified four phenylalanine residues in the active site tunnel in close proximity to the ethylene glycol units of the chloroalkane (CA) substrate (Fig. 1a). We designed four individual point mutants with each of these residues mutated to alanine (Table S1), and we designed a corresponding “bumped” chloroalkane (bCA) substrate with phenylalanine substituted in place of one ethylene glycol unit (Fig. 1b and S2-S4). We hypothesized that the steric bulk of the phenylalanine side chain would inhibit reaction with HaloTag7 while potentially being accommodated by one or more of the Phe-to-Ala mutations.

Using yeast display,^36,40–42^ we screened these four mutants for their ability to react with a BODIPY-linked, bumped chloroalkane substrate (bCA-BDY; data on all four mutants in Fig. S5). We identified HaloTag7 F149A (hereafter referred to as F149A) as the most promising point mutant. To quantify the performance of F149A, we compared the reaction of yeast-displayed HaloTag7 and F149A with bCA-BDY and a non-bumped chloroalkane-BODIPY (CA-BDY) across a range of concentrations from 10 to 250 nM (Fig. 1c). As hypothesized from the bump-and-hole design, yeast cells expressing F149A were up to 11-fold brighter than HaloTag7-expressing cells when labeled with bCA-BDY. Consistent with the structural model, F149A was also less reactive with the non-bumped substrate CA-BDY, so yeast cells expressing HaloTag7 were up to 23-fold brighter when labeled with CA-BDY compared to F149A-expressing cells. Taken together, these data suggested that HaloTag7 and F149A have some mutual orthogonality.

**Figure 1.**
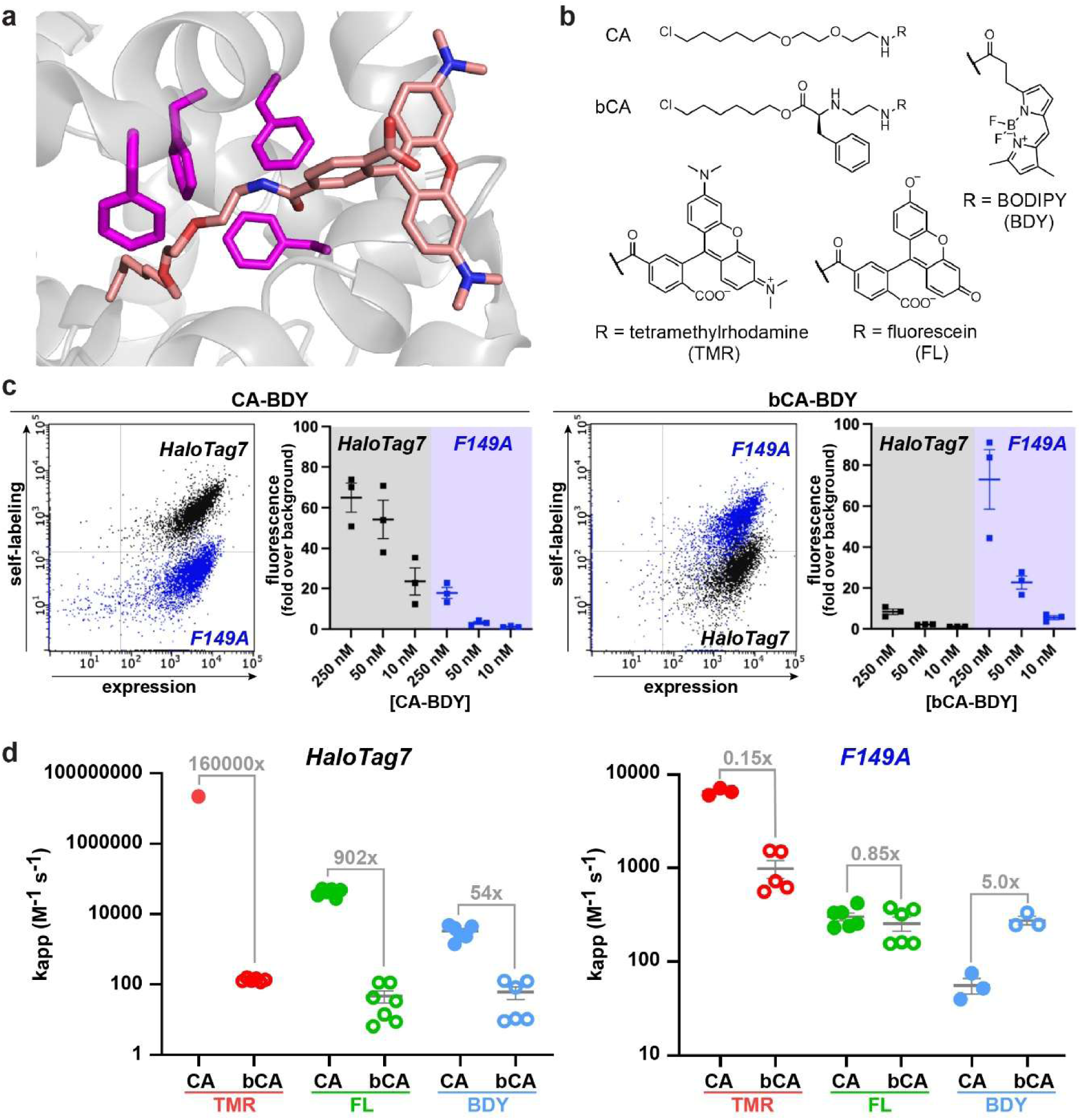
Bump-and-hole engineering of HaloTag7. (a) Four phenylalanine residues (magenta) line the substrate channel of HaloTag7. A covalently bound tetramethylrhodamine HaloTag substrate is shown in salmon (PDB: 6U32).^26^ (b) The canonical HaloTag chloroalkane substrate (CA) has a terminal chloride, a six-carbon chain, and two ethylene glycol units. The new “bumped” chloroalkane substrate (bCA) replaced one of the ethylene glycol units with phenylalanine. This work used CA and bCA substrates attached to BODIPY (BDY), fluorescein (FL), tetramethylrhodamine (TMR), and biotin (Fig. S2-S4). (c) HaloTag7 (black) and the F149A point mutant (blue) were expressed on the yeast surface and treated with 50 nM CA-BDY or bCA-BDY for 15 min. Raw flow cytometry data from yeast display experiments show BDY fluorescence, reporting on self-labeling activity, on the y-axis and immunolabeling for a C-terminal Myc tag, reporting on total expression level, on the x-axis. Compiled cytometry data show the fold increase in green fluorescence compared to non-substrate-treated cells over three biological replicates across multiple substrate concentrations. (d) Kinetics measured using fluorescence polarization revealed k_app_ values for recombinantly expressed and purified HaloTag7 and F149A reacting with six different substrates: CA and bCA, each linked to BDY, FL, or TMR. k_app_ values from three independent trials are shown with mean and standard error of the mean. See Table S2 for kinetics data.

We next recombinantly expressed and purified HaloTag7 and F149A to determine their reaction kinetics with each substrate. We measured self-labeling kinetics using a fluorescence polarization readout, and we measured reaction rates at several enzyme concentrations and a fixed concentration of substrate to produce apparent second-order rate constants.^29^ It has been well-documented that the dye attached to the chloroalkane substrate has a large effect on HaloTag7’s reaction rate,^29^ so we measured kinetics with CA and bCA substrates attached to three different dyes. We found that the extent of orthogonality varied depending on the chosen dye. With BODIPY (BDY), fluorescein (FL) and tetramethylrhodamine (TMR) substrates, the bumped bCA substrate reacted 4.6 to 7.2-fold more rapidly with F149A than with HaloTag7. We also found that the original CA substrate reacted two to three orders of magnitude faster with HaloTag7 than with F149A, regardless of which dye was attached. While the mutual orthogonality was substantial, the apparent rate constants for F149A with bumped substrates were all less than 10^3^ M^-1^ s^-1^. Based on extensive prior work,^22,29,36,43^ this SLP-substrate pair would likely require long labeling times and high substrate concentrations for cellular labeling experiments.

### Directed Evolution to Improve Reaction Kinetics

To enhance reaction rate and orthogonality, we performed directed evolution using yeast display, taking advantage of the platform developed in our previous work evolving HaloTag7 for alternative substrates (Fig. 2a).^36,40–42^ We used error-prone PCR to produce four sub-libraries with different mutational frequencies, and these were pooled into an input library of over 17 million variants of F149A (Table S3). Sequencing confirmed that the input library had an average of 3.8 amino acid mutations relative to F149A (Table S3). When the library was expressed on yeast and treated with bCA-BDY, we observed a broad range of activities (Figs. 2b, S6). Our selection scheme is summarized in Fig. 2c. We performed three rounds of fluorescence-activated cell sorting, collecting the top 1% or less of yeast cells with the highest activity relative to expression level. We alternated substrates between rounds between bCA-FL and bCA-biotin, using streptavidin-AlexaFluor488 to detect self-labeling with bCA-biotin. An excess of nonfluorescent CA substrate was used to bias the sorting towards variants with selectivity for bumped substrates. We increased stringency between rounds by decreasing concentration of bCA-FL or bCA-biotin, decreasing labeling times, and increasing excess CA. We used multiple parallel strategies, pooling between rounds 2 and 3 (Fig. 2c; more details in Supporting Information).

After three rounds of sorting, the enriched output pool had 10-fold brighter fluorescence than F149A when labeled with 60 nM bCA-FL and 300-fold excess of nonfluorescent CA on the surface of yeast (Figs. 2b, S7). Sequencing revealed that no mutations were enriched at position 149, consistent with the constant selective pressure against reaction with non-bumped substrate. One mutation, N272Y, was present in up to 99.8% of the output pool, also indicating strong, directional selection. P142S and F144L mutations were also highly represented, each with some associated mutations: K71R/D187V and I218T co-occurred with P142S, and F91L/G96S co-occurred with F144L (Table S5-S6). We recombinantly expressed and purified F149A variants containing combinations of these mutations and we tested their reaction rates with CA and bCA using fluorescence polarization kinetics assays (Fig. 2d and S8-S9, Table 1). Ultimately, the best-performing mutant was P142S/F144L/F149A/N272Y, which we designated OrthoTag. Covalent labeling of purified OrthoTag with bCA-biotin was confirmed by mass spectrometry (Fig. S10). We measured reaction kinetics of HaloTag7 and OrthoTag with three different dyes, each linked to either the CA or bCA substrate (Fig. 2d, Table S7). As documented previously for HaloTag7,^29^ both enzymes reacted most rapidly with TMR-based substrates, more slowly with FL-based substrates, and most slowly with substrates lacking xanthene dyes.

Notably, OrthoTag had similar reaction rates with CA-TMR and bCA-TMR, suggesting that OrthoTag shares the direct interactions with TMR observed for HaloTag7 and that these interactions are enough to overcome its otherwise reduced reactivity towards CA substrates. By contrast, HaloTag7 has approximately 130,000-fold poorer reaction rate with bCA-TMR than CA-TMR, indicating that the larger size of bCA prevents self-labeling despite highly favorable binding interactions with the TMR group.^29,44^ Compared to HaloTag7, OrthoTag has 109-fold faster labeling with bCA-FL and 140-fold slower labeling with CA-FL (Fig. 2d, Table 1). Between bump-hole engineering and directed evolution, we improved selectivity for the bumped substrate by 17000-fold.

**Table 1.** Kinetic characterization of HaloTag variants. Variants with highly represented mutations from the evolution campaign were recombinantly expressed and kinetics were measured using chloroalkane-fluorescein substrates. Selectivity reflects the ratio of reaction rate for bCA-FL relative to CA-FL, and fold improvement compares bCA selectivity to that of HaloTag7.

|  | Mutations |  |  |  |  |  |  |  |  | k<sub>app</sub> (M<sup>-1</sup>s<sup>-1</sup>) |  | Selectivity for bCA | Fold improvement |
| --- | --- | --- | --- | --- | --- | --- | --- | --- | --- | --- | --- | --- | --- |
|  | F149A | N272Y | F144L | F91L | G96S | P142S | K71R | D187V | I281T | CA-FL | bCA-FL |  |  |
| HaloTag7 |  |  |  |  |  |  |  |  |  | 42500 ± 4020 | 47.2 ± 17.6 | 0.001 | 1 |
| F149A | X |  |  |  |  |  |  |  |  | 297 ± 29 | 252 ± 43 | 0.85 | 850 |
| Variant 1 | X | X | X |  |  |  |  |  |  | 501 ± 31 | 839 ± 91 | 1.7 | 1700 |
| Variant 2 | X | X | X | X | X |  |  |  |  | 372 ± 21 | 659 ± 70 | 1.8 | 1800 |
| Variant 3 | X | X |  |  |  | X |  |  |  | 309 ± 39 | 1660 ± 150 | 5.3 | 5300 |
| Variant 4 | X | X |  |  |  | X | X | X |  | 211 ± 21 | 992 ± 169 | 4.7 | 4700 |
| Variant 5 | X | X |  |  |  | X |  |  | X | 223 ± 9 | 1080 ± 60 | 4.8 | 4800 |
| OrthoTag | X | X | X |  |  | X |  |  |  | 303 ± 11 | 5150 ± 1360 | 17 | 17000 |

**Figure 2.**
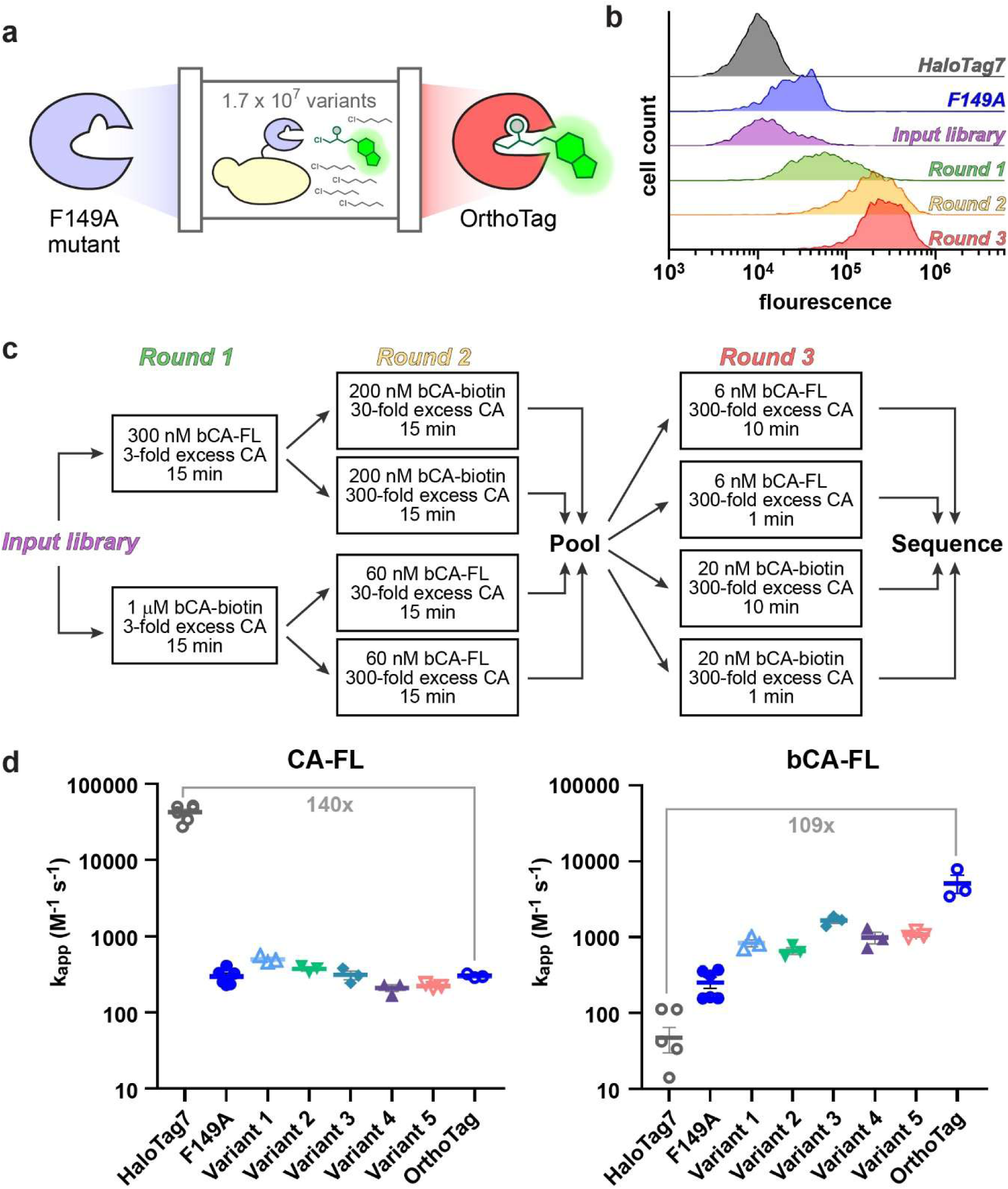
Directed evolution of an orthogonal HaloTag variant. (a) A library of 17 million F149A variants was expressed on yeast and screened using fluorescence-activated cell sorting. (b) Flow cytometry of F149A-expressing yeast, the input library, and the output pools from three rounds of selection. Cells were labeled with 60 nM bCA-FL and 300-fold excess nonfluorescent CA. (c) Summary of selection strategy. Successive rounds alternated bCA substrates, decreased concentration of bCA substrate, decreased incubation time, and increased the excess of non-bumped CA. Additional details in Supporting Information. (d) Kinetics of recombinantly expressed HaloTag7 and variants reacting with CA-FL and bCA-FL. k_app_ values are from at least three independent trials with mean and standard error of the mean. See Fig. S8-S9 and Table S7 for kinetics data.

### Crystal Structures of OrthoTag

We solved the crystal structures of OrthoTag in the *apo* state (PDB ID: 30HW) and in complex with the bumped chloroalkane fluorescein substrate (bCA-FL, PDB ID: 30HV) at 1.40 Å and 2.10 Å resolution, respectively. In the complex, the ligand is covalently attached to Asp106 via nucleophilic substitution of the terminal chloride (key interactions are highlighted in Fig. S11, data collection and refinement statistics in Table S8), in the same manner as the conventional ligand in HaloTag7.^26^ Analysis of the electron density maps (Fig. S11) clearly supports the fluorescein moiety in its open, quinoid form, consistent with the crystallization conditions (pH 6.5), where fluorescein exists in a pH-dependent equilibrium between closed and open states.^45^ The binding sites for TMR in the published HaloTag7 structure and fluorescein in the ligand-bound OrthoTag structure differ, as expected for these different xanthene dyes (Fig. 3a,b).^29^

Comparison of the *apo* and ligand-bound OrthoTag structures reveals that the enzyme adopts an expanded active site architecture even in the absence of substrate. The substrate channel volume in the *apo* structure is 276 Å^3^ and it increases only modestly to 310 Å^3^ upon ligand binding (Fig. 3c).^46^ This represents a large expansion of the substrate channel compared to HaloTag7, which has substrate channel volumes of 162 Å^3^ and 173 Å^3^ for its *apo* and bound states, respectively.^26,46,47^ This small change in cavity volume, along with the RMSD of 0.293 Å between *apo* and *holo* OrthoTag, further supports that the expanded active site is a structural feature of OrthoTag rather than an induced fit upon substrate binding (Fig. S11). Approximately 81 Å^3^ of the substrate channel volume difference between *apo* HaloTag and OrthoTag can be directly accounted for by smaller side chain volumes of mutated residues within the cavity.^48^ The other approximately 33 Å^3^ can be attributed to local backbone shifts (Fig. 3a,b), including the altered structure in residues 142-149 described below. These calculations are approximate because the side chains of S142 and L144 are both partially solvent-exposed and because the electron density of L144 was not completely resolved for its Cy and Cδ atoms (Fig. S11).

Three of the four engineered mutations (P142S, F144L, and F149A, Fig. 3a) localize to a region at the entrance of the substrate channel. This region (T137 to R153) forms a kinked helix in HaloTag7, but in both the *apo* and bCA-FL-bound structures of OrthoTag this region contains an additional turn at S142-L144 (Fig. 3b,d). This structural change, along with decreased steric bulk of the F144L mutation, substantially widens the entrance and upper portion of the substrate channel (Fig. 3d), creating space to accommodate the bulkier bCA ligand as it enters the channel. The substrate channel is likely made more accessible by high sidechain mobility in this region, which is suggested by their degree of solvent exposure and the observations that the side chain electron density for L144 is partially unresolved in both structures and the electron density for the side chain of S142 is partially unresolved in the bCA-FL-bound structure. The wide-open mouth of the substrate channel also allows the dye to be positioned roughly 1 Å closer to the internal active site in OrthoTag, compared to HaloTag7 (Fig. 3c). This difference highlights the less extended, more compact conformation of the bumped substrate within the OrthoTag channel. We measured the relative fluorescence quantum yield (ɸ) of CA-TMR and bCA-TMR, free in aqueous solution and bound to HaloTag and OrthoTag respectively, to determine whether this repositioning affected the photophysical properties of the bound dye (Table S9). CA-TMR showed a modest increase in ɸ when binding to HaloTag, as expected from prior observations.^49,50^ This same increase was not observed for bCA-TMR bound to OrthoTag, a potential result of altered dye-protein interactions.

Despite being the most prevalent mutation in the output library, it was less immediately clear how N272Y contributes to OrthoTag’s enhanced reaction rate and/or selectivity for bumped substrates. Position 272 lies at the base of the active site and corresponds to the catalytic histidine in the parent dehalogenase DhaA, and N272 forms hydrogen bonds with H105 and E130 in *apo* and substrate-bound HaloTag7. In OrthoTag, the Tyr at this position does not make strong polar interactions with H105 or E130. Instead, it establishes new hydrogen bonds with the catalytic residue D106 (Fig. 3e). In the substrate-bound structure, this mutation alters the local conformation of the bound alkane chain, and, overall, it appears to help accommodate the more compact conformation of the bound substrate (Fig. 3c,e).

**Figure 3.**
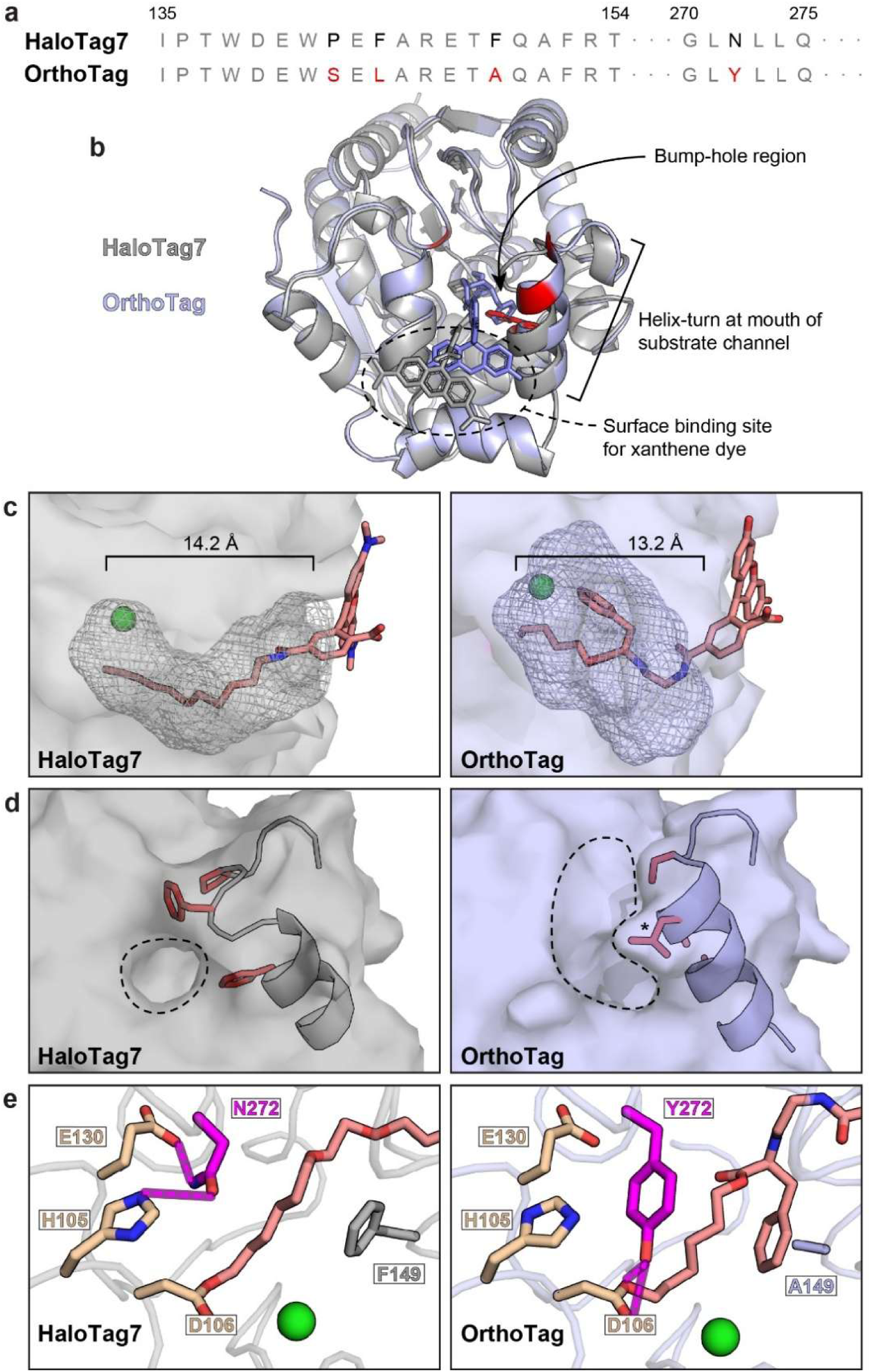
Apo and substrate-bound crystal structures reveal the basis of orthogonality between OrthoTag and HaloTag7. (a) Sequence alignment of HaloTag7 and OrthoTag. Mutated residues are shown in black in the HaloTag7 sequence and red in the OrthoTag sequence. (b) Backbone overlay of substrate-bound HaloTag7 and OrthoTag. Red backbone denotes a mutated residue. The phenyl group of the bCA ligand (blue sticks) occupies approximately the position of the F149 side chain in HaloTag7 (red sticks). (c) Substrate channel volumes for substrate-bound HaloTag7 (PDB: 6U32) and OrthoTag (PDB: 30HV), analyzed using POVME3 (Table S10).^26,46^ apo structures showed similar volume differences. Chloride ions are shown in green. (d) Structures of apo HaloTag7 (PDB: 4KAF) and OrthoTag (PDB: 30HW) with mutated residues shown in red. The dotted circles highlight the widening of the mouth of the substrate channel in OrthoTag. Asterisk denotes L144, for which much of the sidechain is unresolved (Fig. S11c). The conformation of L144 shown here is the rotamer observed in the ligand-bound OrthoTag structure, but it is likely highly mobile, especially for apo OrthoTag. (e) Polar contacts of residue N272 in HaloTag7 and Y272 in OrthoTag (magenta). Hydrogen bonds are shown as dotted magenta lines.

### Multiplexed Labeling in Live Cells

To demonstrate the capabilities of OrthoTag for live cell labeling, we generated two stable HEK293T cell lines. One expressed OrthoTag as a histone 2B fusion (OrthoTag-H2B) localized to the nucleus, and the other expressed HaloTag7 as a TOMM20 fusion (HaloTag-T20) localized to the outer mitochondrial membrane. Western blots confirmed that OrthoTag-H2B and HaloTag-T20 were expressed at roughly equivalent levels (Fig. S16). We chose bCA-TMR and CA-BDY as the substrates, which the kinetics data suggested would maximize two-way orthogonality (Table S7). Using flow cytometry, we observed roughly 10-fold more efficient labeling by bCA-TMR of OrthoTag-H2B-expressing cells compared to HaloTag-T20-expressing cells, with respect to treatment concentration (Fig. 4a). Conversely, we observed roughly 4-fold more efficient labeling by CA-BDY of HaloTag-T20-expressing cells compared to OrthoTag-H2B-expressing cells (Fig. 4a). For both dye-linked substrates applied at mid-nanomolar concentrations, we observed minimal background in HEK293T cells expressing no SLPs and we observed roughly 10-fold brighter labeling in cells expressing the matching SLP compared to cells expressing the other SLP (Fig. S12-14). We also used confocal microscopy to verify that the protein fusions were localized correctly (imaging acquisition parameters in Table S11). OrthoTag-H2B labeling showed clear nuclear localization similar to Hoechst staining (Fig. S17), and HaloTag-T20 labeling showed clear mitochondrial localization similar to MitoTracker™ DeepRed FM staining (Fig. S18). Further, the confocal microscopy confirmed robust labeling and low background in cells at mid-nanomolar concentrations of dye-linked substrates. Importantly, these experiments demonstrated that the bCA substrate was highly permeable, had low background, and were easily washed out at nanomolar concentrations in cultured mammalian cells. To demonstrate that ester hydrolysis was not a major limitation to the performance of the bCA-TMR substrate, we exposed bCA-biotin to freshly prepared cell lysates and monitored its presence by mass spectrometry (Fig. S15). This assay is a stringent test for hydrolytic stability.^51,52^ While a control chloroalkane-tagged peptide was completely undetectable after 1 hour, we saw no evidence of degradation of bCA-biotin for up to 3 hours.

To perform multiplexed labeling in live cells, we generated a stable HEK293T cell line expressing both OrthoTag-H2B and HaloTag-T20. Western blots confirmed that OrthoTag-H2B and HaloTag-T20 were expressed at similar levels in this cell line (Fig. S16). We treated these cells simultaneously with 100 nM bCA-TMR and CA-BDY and we used confocal microscopy to visualize their subcellular localization (Fig. 4b and S19). We observed clear localization of bCA-TMR to nuclei and localization of CA-BDY to mitochondria, with minimal cross-labeling. Quantitative colocalization analysis showed correlation between the nuclear stain and bCA-TMR signals and mild anti-correlation between bCA-TMR and CA-BDY signals, indicating orthogonal labeling (Fig. 4c). We further verified in-cell target engagement through an in-gel fluorescence assay. Using optimized treatment conditions (200 nM bCA-TMR added first and 50 nM CA-BDY added after 30 min), we observed clear bands of in-gel fluorescence at the expected positions of the OrthoTag-H2B and HaloTag-T20 fusion proteins with minimal, if any, cross-labeling observed (Fig. S20).

**Figure 4.**
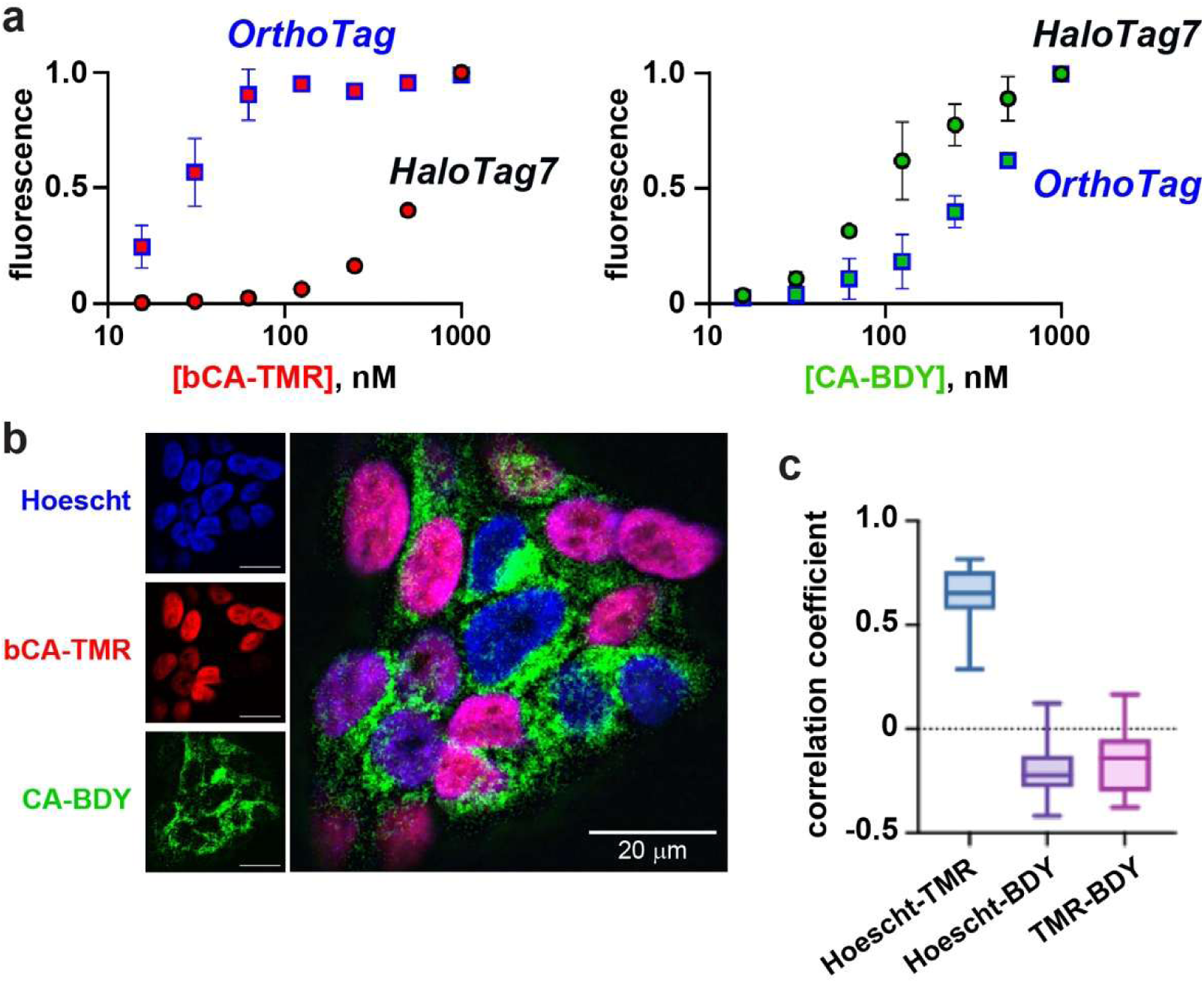
Multiplexed labeling in mammalian cells. a) Flow cytometry of HEK293T cells expressing either HaloTag-T20 or OrthoTag-H2B and labeled with different concentrations of bCA-TMR (left) or CA-BDY (right). Fluorescence was normalized to maximum fluorescence and background fluorescence for each experiment, and data are the averages and standard errors of the mean for three biological replicates. b) Representative confocal fluorescence microscopy image of HEK293T cells expressing both HaloTag-T20 and OrthoTag-H2B and labeled with 100 nM CA-BDY and 100 nM bCA-TMR for 1 h before washing, fixing, and staining with Hoechst 33342. c) Correlation coefficients for the Hoechst, TMR, and BDY channels from confocal fluorescence microscopy (n = 20 cells). The line shows the median value, boxes show the middle quartiles, and whiskers show the minimum and maximum values in each dataset. Coefficients range from -1 to 1, with -1 representing perfect anti-localization and 1 representing perfect colocalization. Coefficients were calculated using CellProfiler.^53^

## Discussion

Fluorescent proteins remain a popular tool in microscopy and as components of protein-based biosensors due to their genetic targetability and ease of use. However, applications are restricted by their limited brightness, poor photostability, and maturation times.^5,6^ HaloTag can be paired with optimized synthetic dyes to overcome many of these issues, unlocking advanced applications including robust single-molecule microscopy and tunable far-red/near-IR biosensors.^20,25,26,54^ Additionally, HaloTag is modular and can be used for the precise localization of other functional compounds such as photocatalysts, affinity handles, reactive metabolites, and drugs.^7–19^ In many cases, fluorescent proteins or luminescent reporter enzymes are paired with HaloTag for the design of FRET/BRET-based sensors.^20,24–26^ In all these applications, other SLPs such as SNAP-tag and CLIP-tag have seen limited use compared to HaloTag due to their slower reaction kinetics, reduced substrate permeability, poorer washout, and interactions with endogenous proteins.^2,3,22,29,31^

Using the “bump and hole” method for the rational design of orthogonal enzymes,^37–39^ we produced a HaloTag7 F149A point mutant which reacts selectively with a sterically bulky chloroalkane substrate (bCA). The bump-and-hole mechanism was shown structurally: when the substrate-bound OrthoTag structure is overlayed with the structure of HaloTag7, the phenyl group of the bumped ligand occupies almost the same space occupied by the phenyl group of F149 in HaloTag7 (Fig. 3b,e). Using yeast display, we further improved reaction rate by 20-fold while maintaining a low reaction rate with the original chloroalkane substrate (CA). OrthoTag expresses robustly in bacterial, yeast, and mammalian cells, and our observations to date suggest the bCA substrate is comparable to the linear chloroalkane with respect to cell permeability and washout, ensuring low background. We demonstrated that the combination of HaloTag7 and OrthoTag enables rapid, simultaneous, orthogonal labeling in living cells at substrate concentrations at and below 100 nM.

Prior HaloTag engineering efforts have primarily focused on tailoring dye-protein interactions at the enzyme’s surface. Modifications to the chloroalkane linker and remodeling of the deeper HaloTag active site tunnel remain relatively underexplored, besides a single recent report of an alternative chloroalkane substrate which improved conjugation to neuronally expressed HaloTag7.^14^ Crystal structures of *apo* and substrate-bound OrthoTag revealed a large widening of the upper portion of the active site tunnel, as well as a more subtle rearrangement at the bottom of the tunnel near the catalytic D106. The specific mutations which support this rearrangement are informative for continued efforts in HaloTag engineering.^15,16,20,24–26,34,36,44^ N272Y is a particularly important mutation, as it occupies the position of the catalytic histidine in the parent dehalogenase. H272 was initially mutated to Phe in the earliest iteration of HaloTag design, a critical mutation which removed the ester hydrolysis activity of the parent enzyme to produce a self-labeling protein.^27^ Asn at position 272 was subsequently shown to improve stability without restoring undesirable hydrolytic activity.^28^ Subsequent design and evolution efforts have not altered N272,^15,16,20,24–26,34,36,44^ making the N272Y mutation in OrthoTag unique. The roles of Phe and Asn residues at this position were recently explored in the context of dehalogenase activity,^55^ but Tyr was not explored. Our structures suggest that Y272 may remodel the region surrounding the catalytic D106 residue, perhaps acting as a gatekeeper to selectively promote reactions with bumped substrates over linear substrates. Since our kinetic data do not allow decomposing k_app_ into k_cat_ and k_M_, more extensive pre-steady-state kinetics would be required to elucidate conclusively the role of Y272 in substrate recognition and catalysis.^29^ The mutations P142S and F144L both support widening of the mouth of the active site tunnel. Previous work adapting HaloTag7 for use with styrylpyridium and benzothiadiazole dyes identified F144 as a position which modulates reaction rate and dye microenvironment once attached.^36,47,56,57^ By contrast, as far as we are aware, this is the first example of a desirable mutation at the P142 position. F149A serves to widen the central portion of the substrate channel, and thus it is likely only desirable for substrates with increased steric bulk closer to the terminal alkane. More broadly, our work demonstrates that the HaloTag active site can be dramatically remodeled, for instance making the substrate channel 70% larger, while maintaining robust expression and activity. This work provides a clear basis for developing SLPs which can accommodate larger substrates.^58,59^

OrthoTag maintains a large degree of sequence and structural homology to HaloTag7, and its bumped substrate closely resembles the preferred HaloTag substrate. Thus, we anticipate that it will be readily substituted into existing HaloTag-based platforms to allow multiplexing with minimal re-optimization. Such applications would be more complex to develop with unrelated SLPs such as SNAP-tag2.^31^ Observed labeling efficiencies in single-expressing and dual-expressing cell lines inform the selection of substrate-dye conjugates and inform conditions for maximizing orthogonality. For example, CA-rhodamine substrates should be avoided, because these substrates retain higher activity towards OrthoTag due to its intrinsic affinity for rhodamines. Additionally, the bumped substrate can be added at a higher concentration and allowed to react for 30 min before adding the linear substrate, as HaloTag7 has very low labeling efficiency for a range of bCA-dyes. In the future, OrthoTag could potentially be applied for orthogonal labeling with multiple fluorogenic dyes in a no-wash format,^36^ orthogonal localization of two different neurotransmitters to two different neuronal subtypes using the DART method,^9,14^ orthogonal molecular recorders to measure correlations between two cellular events in live animals,^15,16^ orthogonal photoproximity labeling at multiple sites in the cell using HaloMap,^11,17^ or temporal and spatial control of multiple reactive metabolites using multiplexed T-REX.^7^ The design and evolution methods described here could also be directly applied to develop a third, mutually orthogonal HaloTag variant. With this additional SLP, and/or with optimization of SNAP-tag2 for use with these biotechnology methods, there is potential to multiplex these experiments to address three or more targets or processes in the cell.^31,60,61^

## Acknowledgments

This work was supported by NSF MCB 2530334 to J.A.K and NIH 5F32GM139371 to B.J.L. We thank the staff at Diamond Light Source UK for beam time at beamlines I03 and I04 (proposals MX-31353 and MX-38144). P.R. thanks the Wellcome Trust (227298/Z/23/Z) and C.J.S. for continuous support and use of instruments. A.T.S acknowledges funding from the EPSRC (EPW524311/1) and Diamond Light Source.

## Data availability

The atomic coordinates and structure factors are deposited in the PDB with the following accession codes: 30HW (*apo* Orthotag) and 30HV (bCA-FL-bound Orthotag).

## Methods

### Substrate Synthesis and Purification

Synthesis protocols and characterization data for chloroalkane substrates are provided in the Supplementary Information (Fig. S2-S4).

### Yeast Culture and Molecular Biology

HaloTag7, codon-optimized for *S. cerevisiae*, was cloned into yeast display vector pCTcon2 (originally produced by Dane Wittrup,^62^ Addgene #41843) as previously described.^36^ RJY100 *S. cerevisiae* were used for all experiments.^63,64^ Yeast stocks were stored at -80 °C in 15% glycerol solutions.

DNA sequences encoding F144A, F149A, F152A, and F168A point mutants of HaloTag7 were synthesized by Twist Bioscience with homologous overhangs (Table S1), amplified using primers **P3** and **P4** (Table S12), and cloned into pCTcon2 using Gibson Assembly® Master Mix (New England Biolabs).

Tryptophan auxotrophic RJY100 *S. cerevisiae* transformed with pCTcon2 were selected using a Trp biosynthesis marker and cultured according to published protocols.^36,40^ Briefly, *S. cerevisiae* were grown in tryptophan dropout synthetic dextrose media (SD -Trp, pH 4.5) at 30 °C. To perform a display experiment,^36^ *S. cerevisiae* cultures were diluted to 1 x 10^7^ cells/mL the day prior and grown to mid-log phase at 30 °C. To induce expression of Aga1p and the Aga2p-HA-HaloTag-Myc construct, cells were pelleted by centrifugation and resuspended in tryptophan dropout synthetic galactose media (SG -Trp, pH 6.5) at a concentration of 1 x 10^7^ cells/mL and left to induce overnight at room temperature.

The morning of the experiment, 2 x 10^6^ cells were collected from each induced culture and pelleted before washing with 50 µL of phosphate-buffered saline (PBS). HaloTag labeling was performed with 50 µL of a solution of the appropriate substrate in PBS at room temperature on a rotator. After labeling, the solution was aspirated, and cells were washed again three times. Primary antibody labeling was performed in 50 µL of room temperature PBSA (PBS + 0.1% w/v bovine serum albumin) with either rabbit anti-HA or mouse anti-Myc antibody diluted 1:2000, for at least 30 min with rotation, before removing and incubating for 15 min with goat anti-rabbit Alexa Fluor (AF) 547 or goat anti-mouse AF-488 on ice. Making sure to maintain ice-cold conditions, cells were washed once with PBSA before pelleting and resuspending in 200 µL PBSA for flow cytometric analysis.

### Library Construction, Cell Sorting, and Directed Evolution

Libraries of HaloTag mutants were constructed according to established protocols.^36,40,42,65^ Standard PCR (Table S13) was performed on DNA encoding HaloTag7 F149A with the addition of either 2 or 10 µM 8-oxo-2′-deoxyguanosine-5′-triphosphate and 2′-deoxy-P-nucleoside-5′-triphosphate (Jena Bioscience) for 10 or 20 cycles to induce random mutations (Table S3). Amplicons were purified using the QIAquick® Gel Extraction Kit (Qiagen) and re-amplified for 35 cycles of standard PCR. Inserts were transformed into electrocompetent RJY100 *S. cerevisiae* by electroporation according to prior published protocols with linearized pCTcon2 vector.^36,40^ Library size was estimated by plating dilutions,^36^ average mutation frequency was determined by single colony Sanger sequencing analysis (n ≥ 20) (Azenta Life Sciences GENEWIZ), and percent activity relative to parent was characterized by labeling with bCA followed by flow cytometry (Fig. S6, Table S3). Because all sub-libraries were deemed to have sufficient activity, they were pooled at 10x estimated diversity and used for subsequent directed evolution experiments.

For each round of sorting, a number of cells 10x the estimated diversity of the input library was prepared as described above, with volumes scaled linearly with respect to the number of cells. Labeling conditions were changed between rounds (Table S4). The top 0.1 to 1% of brightest cells were collected on a Bio-Rad S3e Cell Sorter by drawing a sorting gate above the main population accounting for activity (enzyme self-labeling, green fluorescence) and full-length expression (anti-Myc immunostaining, red fluorescence). Sorted cells were recovered in SD -Trp media with 1% penicillin/streptomycin and allowed to grow at 30 °C until solutions appeared cloudy, before flow cytometry characterization and storage as described above. Output populations from each round were pooled and used for the next round of sorting. After the third round, the output pool was only ∼6 x 10^3^ highly active mutants. To sequence hits, yeast cells were lysed and DNA collected by Zymoprep Yeast Plasmid Miniprep Kit (Zymo Research) before PCR amplification of the HaloTag gene using standard conditions and sequencing (Plasmidsaurus Premium PCR Sequencing).

### Mammalian Cell Culture, Flow Cytometry, and Stable Cell Line Generation

All cell culture media and reagents were purchased from ThermoFisher Scientific unless otherwise specified. Lentiviral vectors encoding HaloTag7-TOMM20, or OrthoTag-H2B, or a fusion of the two with a self-cleaving P2A linker were produced by VectorBuilder. Lentiviral packaging vectors psPAX2 (Addgene #12260) and VSV.G/pMD2.G (Addgene # 2259) were used. Lentivirus was packaged in-house following established protocols.^66,67^

HEK293T cells were cultured in Dulbecco’s Modified Eagle Medium (DMEM) with 4.5 g/L D-Glucose, 584 mg/L L-Glutamine, 110 mg/L sodium pyruvate, and 10% v/v fetal bovine serum (FBS) (Neuromics). 5 µg/mL of blasticidin S HCl was added for selection of stable cell lines (Fig. S21), and 2 µg/mL blasticidin S HCl was used thereafter to maintain selection. Cell lines were tested for mycoplasma using a Universal Mycoplasma Detection Kit (ATCC).

To prepare cells for flow cytometry, 2 x 10^4^ cells were plated in each well of a tissue-culture treated 96-well plate 24 hours before an experiment in 200 µL of full media. The day of the experiment, 100 µL of serial dilutions of dye-linked substrates were prepared in Opti-MEM reduced serum media without phenol red and added to the cells after aspirating culture media. After incubating at 37 °C for one hour, three washes were performed with 100 µL of Opti-MEM, and then cells were detached in 40 µL of 0.05% trypsin-EDTA w/o phenol red for 5 min at 37 °C. Finally, cells were resuspended with 160 µL of PBS and then analyzed on an Agilent Novocyte Quanteon flow cytometer.

### Confocal Microscopy

Fluorescence confocal microscopy was performed with a Leica FLIM SP8 on fixed HEK293T cells. Two days before the experiment, 5 x 10^8^ cells were plated in individual wells of a 6-well plate containing three 12 mm cover glasses (Paul Marienfeld GmbH and Co.) and 2 mL of full culture media. Cells were allowed to grow overnight before treating and washing as described in the previous section, using 1 mL of treatment/washing solution for each step to ensure even coverage and being careful to avoid disturbing adhered cells.

To fix, cells were washed three times with 1 mL PBS before incubating with 2 mL of 4% paraformaldehyde in PBS for 15 min at room temperature. Cells were washed three times with PBS again and incubated for 5 min at room temperature with 1 mL of PBS and 0.4 µL of 10 mg/mL Hoechst 33342 trihydrochloride trihydrate in water (Life Technologies). A final three washes were performed, and cover slips were gently dried with a sterile wipe before placing cell-side down on a scant drop of ProLong™ Gold Antifade Mountant. Mounted slides were allowed to dry overnight at room temperature before imaging the next day using either the 40x or 63x water immersion objective of the Leica FLIM SP8. Dry slides were stored at 2 to 8 °C for short-term use.

For mitochondrial staining, MitoTracker™ DeepRed FM was used following manufacturer protocols after washing out excess chloroalkane substrate and prior to fixing cells.

### Protein purification for crystallography studies

For crystallization studies, OrthoTag containing an N-terminal hexa-histidine tag in a pET vector was transformed into *Escherichia coli* BL21 (DE3) cells (New England Biolabs, UK) and grown on 2xYT agar plates supplemented with ampicillin (100 µg/mL). A single colony was picked to inoculate 100 mL of 2xYT broth supplemented with ampicillin and grown at 37 °C overnight. 8 mL of this culture was added to flasks containing 800 mL of 2xYT supplemented with ampicillin and grown at 37 °C until an OD_600_ (optical density at 600 nm) of 0.65 was reached. Protein expression was induced by addition of 0.5 mM IPTG, and cultures were grown overnight at 18 °C. Cells were harvested by centrifugation (12, 230 x *g*, 15 min, 4 °C) and stored at -80 °C. Frozen cell pellet was resuspended (1:4 w/v) in buffer A (50 mM Tris pH 7.4, 200 mM NaCl, 5 mM imidazole, 1 mM 2-mercaptoethanol) supplemented with deoxyribonuclease I (10 µg/mL), phenylmethylsulfonyl fluoride (10 µg/mL), and lysozyme (0.2 mg/mL) by stirring at room temperature for 20 minutes. Cells were lysed using a high-pressure cell disruptor (25 kPSI, 4 °C) and the lysate was clarified by centrifugation (58, 540 x *g*, 30 min, 6 °C). The supernatant was filtered (0.45 µM, Sarstedt, Germany) and loaded onto a Ni-NTA column (5 mL HisTrap HP, Cytiva) pre-equilibrated with buffer A. The column was washed with 15 column volumes of buffer A prior to elution using a linear gradient from buffer A to buffer B (50 mM Tris pH 7.4, 200 mM NaCl, 500 mM imidazole, 1 mM 2-mercaptoethanol) over 10 column volumes. Fractions containing OrthoTag were identified using SDS-polyacrylamide gel electrophoresis (PAGE) and combined. Removal of the N-terminal hexa-histidine affinity tag was achieved by incubating the protein with TEV protease (1:100 w/w) in dialysis tubing (8000 Da MWCO, Thermofisher Scientific, UK) overnight at 4 °C in buffer C (50 mM Tris pH 7.4, 200 mM NaCl, 1 mM 2-mercaptoethanol). Untagged OrthoTag was then purified by passing over a Ni-NTA column pre-equilibrated with buffer A, with unbound fractions collected. OrthoTag was then concentrated by centrifugation (10,000 MWCO, 3148 x *g*, 4 °C, Amicon Ultra) and further purified using size exclusion chromatography (Superdex S75 16:600, Cytiva, UK, column equilibrated with buffer D: 25 mM Tris, 100 mM NaCl, 1 mM 2-mercaptoethanol). OrthoTag-containing fractions were combined and concentrated by centrifugation to 12 mg/mL, aliquoted, flash frozen in liquid nitrogen, and finally stored at -80 °C.

### Crystallization of OrthoTag, X-ray diffraction data collection, and data processing

For crystallisation of bCA-FL-bound OrthoTag, 5.4 µL of bCA-FL (20 mM stock in DMSO) was added to 352 µL of 100 µM OrthoTag (diluted in buffer D), and incubated together overnight at 4 °C under protection from light, with formation of the covalent complex confirmed by LC-MS (*apo* OrthoTag observed mass: 33538 Da, bCA-FL-bound OrthoTag observed mass; 34186 Da, with > 95% labelling efficiency). The complex was subsequently concentrated to an approximate final concentration of 12 mg/mL and mixed in a 2:1 ratio with mother liquor (0.2 M NaCl, 2.0 M ammonium sulfate, 0.1 M sodium cacodylate pH 6.5). Crystals were grown at 10 °C using sitting drop vapour diffusion (300 nL final drop volume). For crystallisation of *apo* OrthoTag, 12 mg/mL OrthoTag was mixed in a 2:1 ratio with mother liquor (0.2 M sodium acetate trihydrate, 0.1 M sodium cacodylate pH 6.5, 30 % w/v PEG-8000) and dispensed as described above. Crystals were grown at 25 °C. For both conditions, crystals were grown for 3 days, before being cryoprotected with 20% glycerol (v/v). Crystals were then immediately harvested with a nylon loop and plunged into liquid nitrogen. Single-crystal diffraction data was collected at 100K at beamlines I03 (*apo* OrthoTag) and I04 (bCA-FL-bound OrthoTag) at the Diamond Light Source. Data was processed using the Xia2 Dials pipeline.^68^ The structure was solved using isomorphous molecular replacement using Phaser,^69^ with PDB 6Y7A used as a search model.^29^ Refinement was iteratively performed using PHENIX.refine with model building in Coot.^70,71^ Ligand restraints were obtained using the Grade2 Web Server.^72^ The final coordinates and structure factors were deposited in the PDB using the following accession codes: 30HW (*apo* OrthoTag) and 30HV (bCA-FL-bound OrthoTag). Data collection and refinement statistics are given in Table S8.

## Notes

### Competing Interest Statement

The authors have declared no competing interest.

### Summary of Updates

updated with additional data and analysis, including more robust demonstration of orthogonal labeling in cells

