## supplemental data for "Design and Evolution of an Orthogonal HaloTag for Multiplexed Labeling in Cells"

**Supplementary Information**

### Analysis of Physicochemical Properties of Labeling Protein Substrates

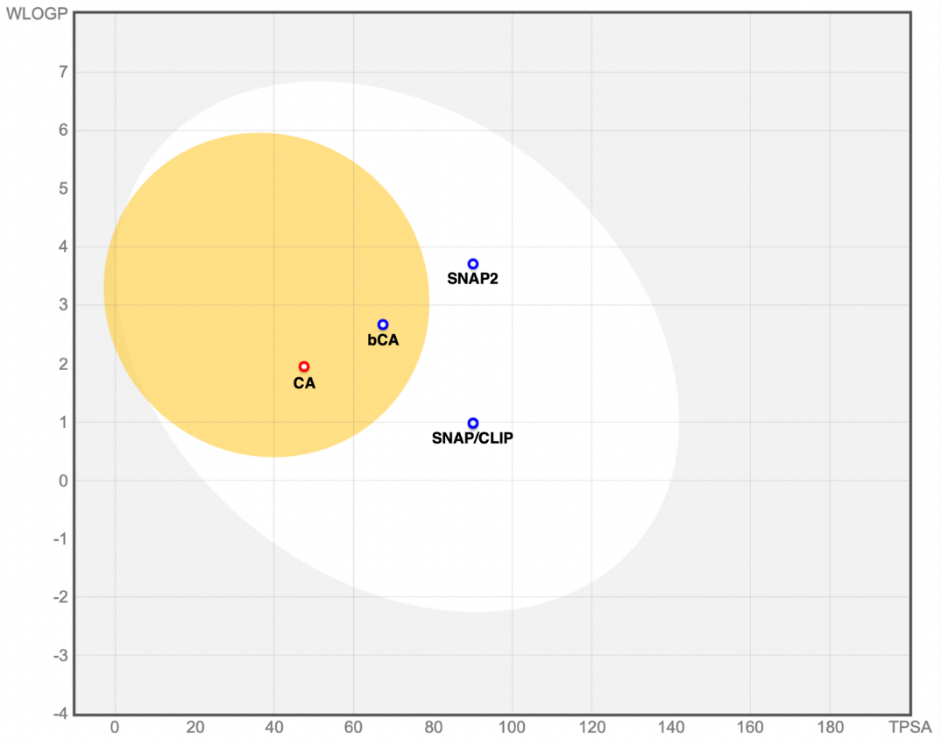

#### **Supplemental Figure 1. Plot comparing physicochemical properties of chloroalkane substrates CA and bCA to substrates for SNAP-tag and CLIP-tag.** The x-axis shows total polar surface area and the y-axis shows WlogP. Compounds which sit in the white circle are predicted to be absorbed through the GI tract, while compounds in the yellow circle are predicted to be blood-brain barrier penetrant. Blue dots indicate compounds that are predicted to be actively effluxed by P-glycoprotein, red dots indicate compounds that are not. Substrates were plotted as acetylated amines to compare their properties in the absence of conjugated fluorescent dyes. This plot was generated using SwissADME.

### Compound Synthesis and Characterization

Dye-linked HaloTag7 substrates were synthesized following prior protocols.^1,2^ Fluorescent dyes were purchased from LumiProbe as NHS esters for conjugation to chloroalkane-amines. Biotin-NHS ester was purchased from Thermo Fisher Scientific. The pure 6-isomer of tetramethyl rhodamine and fluorescein were used. Amino acids were purchased from Chem-Impex. All other synthetic reagents were purchased from Millipore Sigma, TCI, or Thermo Fisher.

#### **Supplemental Scheme 1. Synthesis of the bumped chloroalkane substrate.**

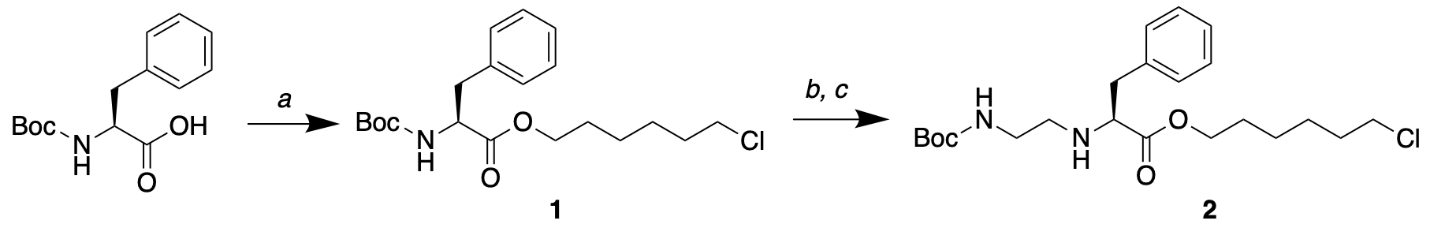

*a. dicyclohexylcarbodiimide, dimethylaminopyridine, 6-chlorohexanol, DCM, 18 hours
b. 4M HCl/Dioxanes, 30 minutes
c. N-Boc-2-aminoacetaldehyde, acetic acid, NaCNBH_3_, MeOH, 36 hours*

**Synthesis of 1.** *N*-Boc-L-phenylalanine (0.126 mmol, 35.2 mg), 6-chlorohexanol (0.189 mmol, 25.82 mg), dicyclohexylcarbodiimide (0.139 mmol, 28.68 mg), and dimethylaminopyridine (0.0126 mmol, 1.54) were dissolved in ice cold dichloromethane (1 mL) and stirred for 5 min. The solution was removed from the ice bath and allowed to stir at room temperature overnight before concentrating *in vacuo* to yield a crude oil. The crude oil was purified by silica gel chromatography, and the target compound was isolated as a white solid (37.25 mg, 74.5% yield).

**Synthesis of 2.** Compound **1** (0.261 mmol, 100 mg) was dissolved in HCl/dioxane (4 M, 1 mL) and stirred at room temperature for 30 min before concentrating *in vacuo* to yield a crude solid. The crude solid was dissolved in methanol (2 mL) before adding *N*-Boc-2-aminoacetaldehyde (0.522 mmol, 83 mg), acetic acid (0.261 mmol, 15 µL), and sodium cyanoborohydride (0.391 mmol, 25 mg). The solution was stirred at room temperature for 36 hours and then diluted in water and basified to pH 12 before extracting with ethyl acetate. The organic layer was dried and concentrated *in vacuo* and purified by silica gel chromatography to yield the target compound as a pale yellow oil (70 mg, 64.8% yield). ^1^H NMR (400 MHz, CDCl_3_) δ 1.28 (p, 2H, J = 7.1 Hz), 1.40 to 1.48 (m, 12H), 1.55 to 1.59 (m, 2H), 1.77 (p, 2H, J = 6.90 Hz), 2.56 to 2.62 (m, 1H), 2.77 (p, 1H, J = 5.90 Hz), 2.94 to 2.96 (m, 2H), 3.07 to 3.21 (m, 2H), 3.50 (t, 1H, J = 7.0 Hz), 3.55 (t, 2H, J = 6.7 Hz), 4.06 (t, 2H, J = 6.6 Hz), 4.87 (s, 1H), 7.19 to 7.33 (m, 5H). ^13^C NMR (400 MHz, CDCl_3_) δ 25.17, 26.45, 28.38, 28.43, 32.39, 39.77, 40.09, 44.92, 47.32, 62.50, 64.70, 79.11, 128.78, 128.45, 129.19, 137.27, 156.02, 174.61. LC-MS (ESI+) exact mass calculated for C_22_H_36_ClN_2_O_4_ (MH^+^) 427.23, observed 427.2822.

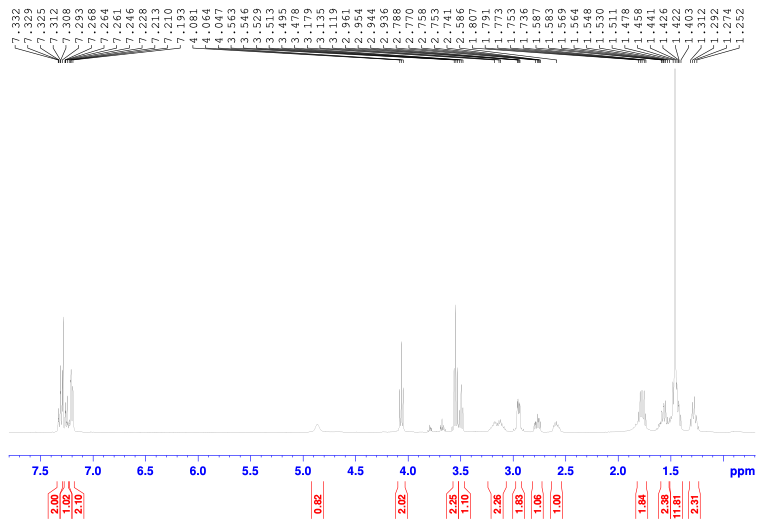

#### **Supplemental Figure 2.** **^1^H NMR spectrum of 2 in CDCl_3_.**

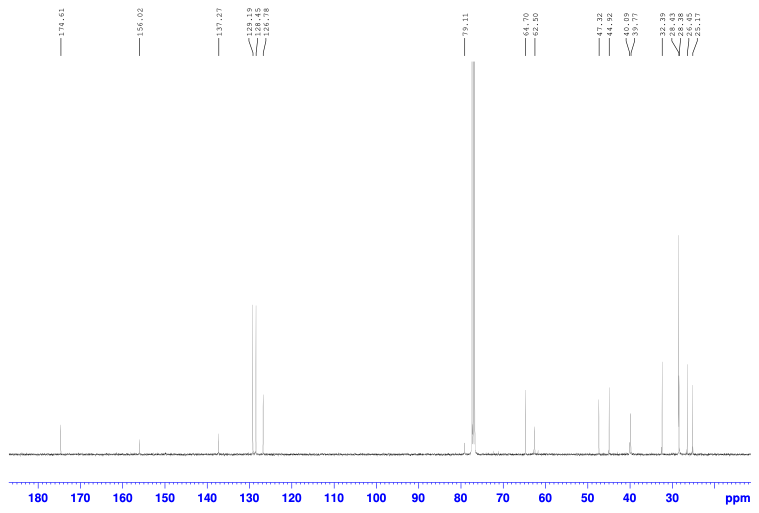

#### **Supplemental Figure 3.** **^13^C NMR spectrum of 2 in CDCl_3_.**

#### **Supplemental Scheme 2.** **Conjugation of dye to chloroalkane substrates**

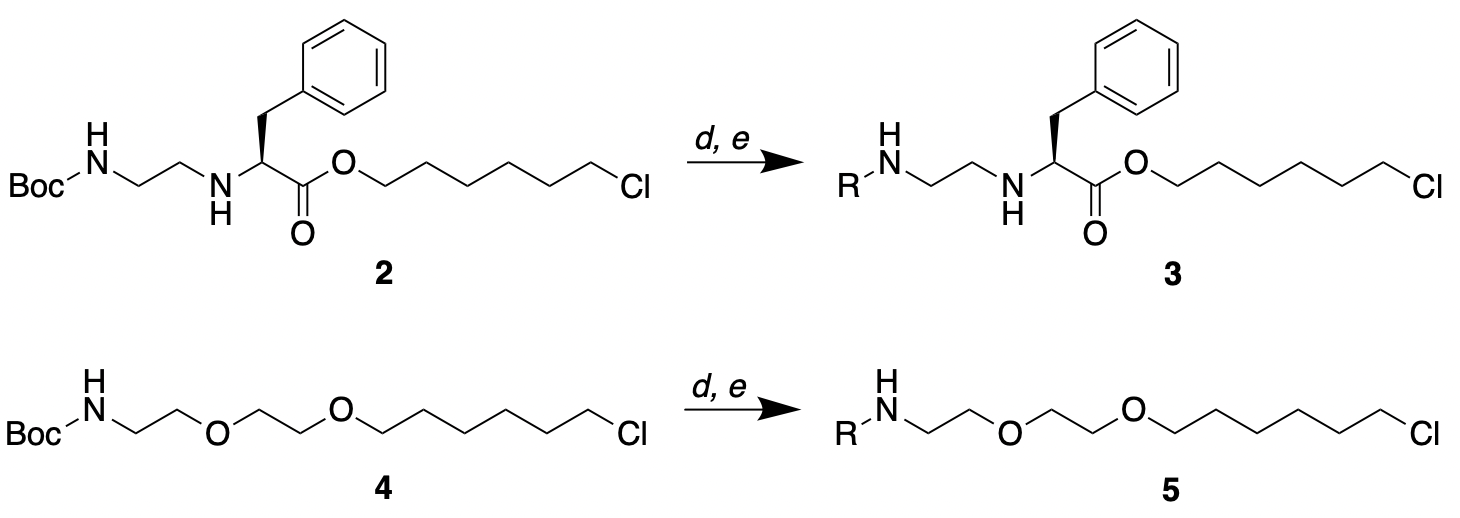

#### d. 4M HCl/Dioxanes, 30 minutes

#### e. R = tetramethylrhodamine/BODIPY-FL/fluorescein/biotin-PEG4 NHS ester, diisopropylethylamine, DMF, 24 hours

**Synthesis of compounds 3 and 5.** Compound **2** or **4** (0.022 mmol) was dissolved in HCl/dioxane (4 M, 0.5 mL) and stirred at room temperature for 30 min before concentrating *in vacuo* to yield a crude solid. The crude solid was dissolved in DMF (300 μL) before adding dye NHS ester (0.015 mmol) and diisopropylethylamine (0.075 mmol, 13 μL) and rotating overnight at room temperature. The solution was diluted in 50:50 acetonitrile:water and purified by reverse-phase HPLC to yield the target compound as a magenta (R = TMR), orange (R = FL, R = BDY), or white solid (R = Biotin-PEG4).

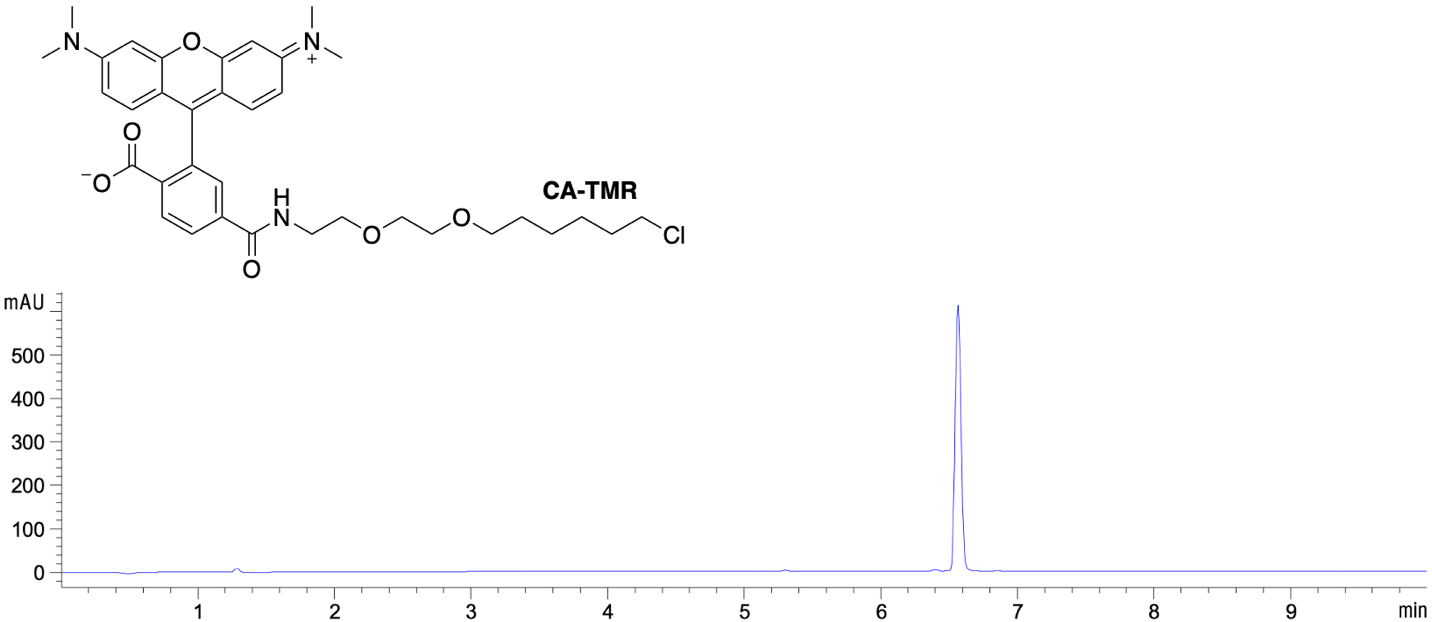

HPLC characterization of CA-TMR at 545 nm. LC-MS (ESI+) exact mass calculated for C_35_H_43_ClN_3_O_6_ (MH^+^) 636.28, found 636.3035.

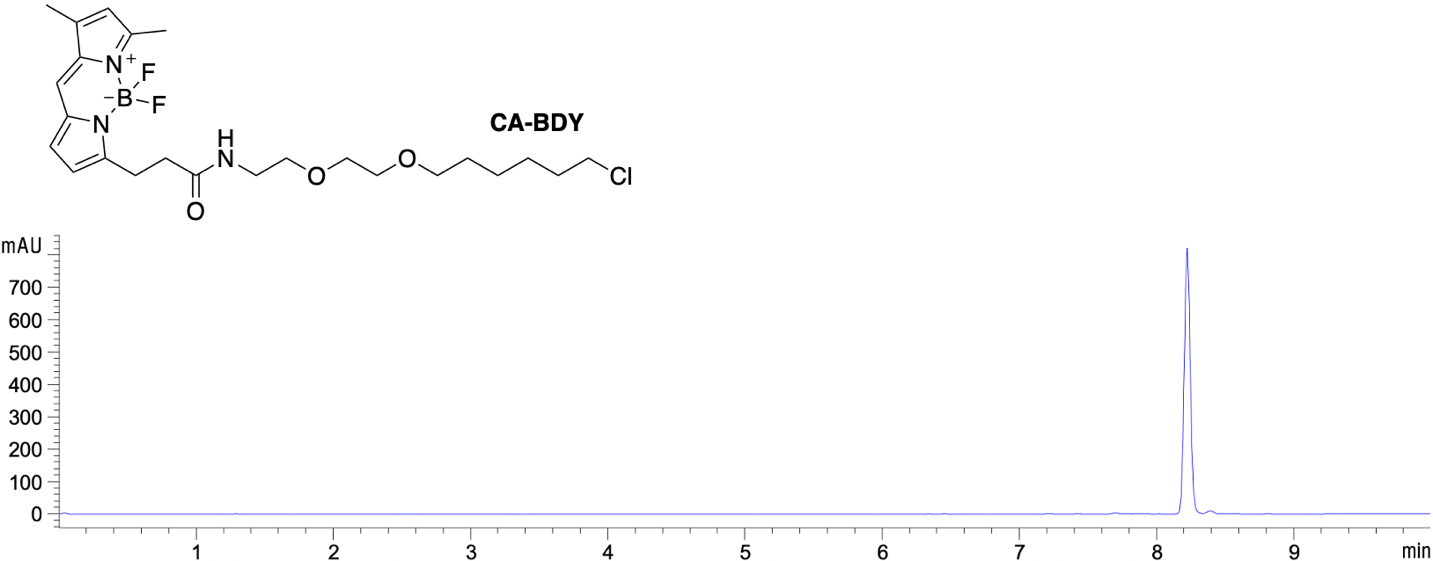

HPLC characterization of CA-BDY at 495 nm. LC-MS (ESI+) exact mass calculated for C_24_H_36_BClF_2_N_3_O_3_ (MH^+^) 498.24, found 498.2560.

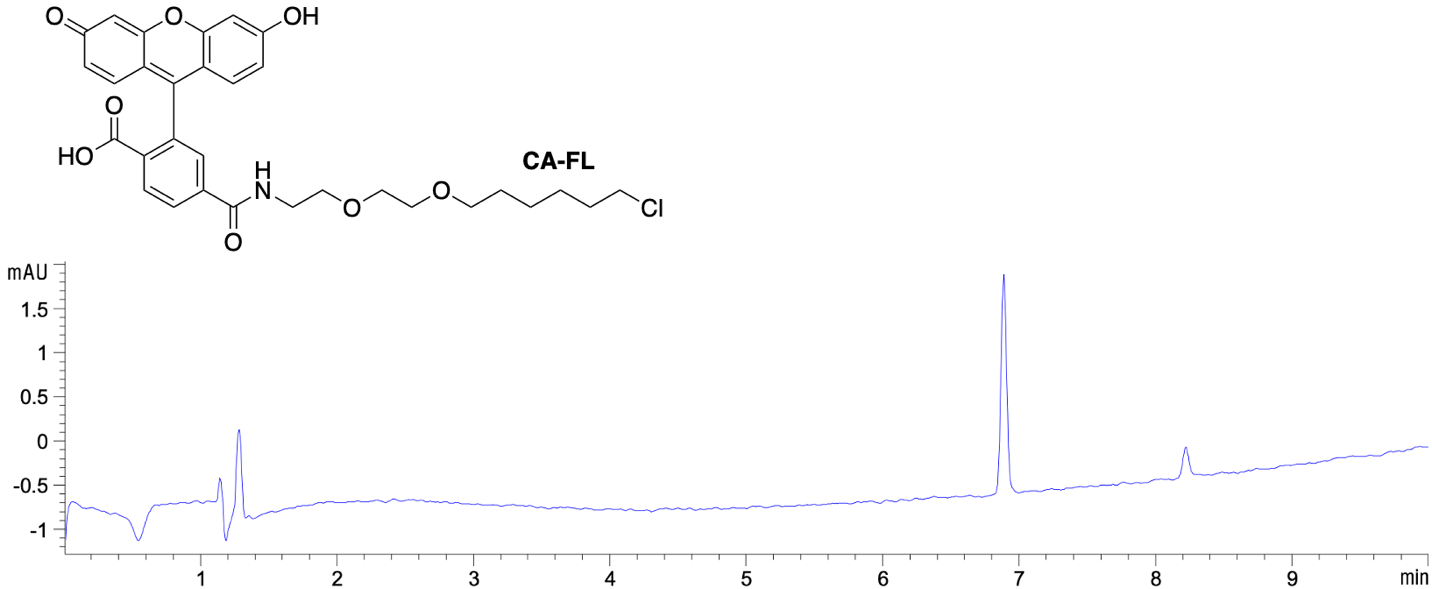

HPLC characterization of CA-FL at 495 nm. LC-MS (ESI+) exact mass calculated for C_31_H_33_ClNO_8_ (MH^+^) 582.18, found 582.1906.

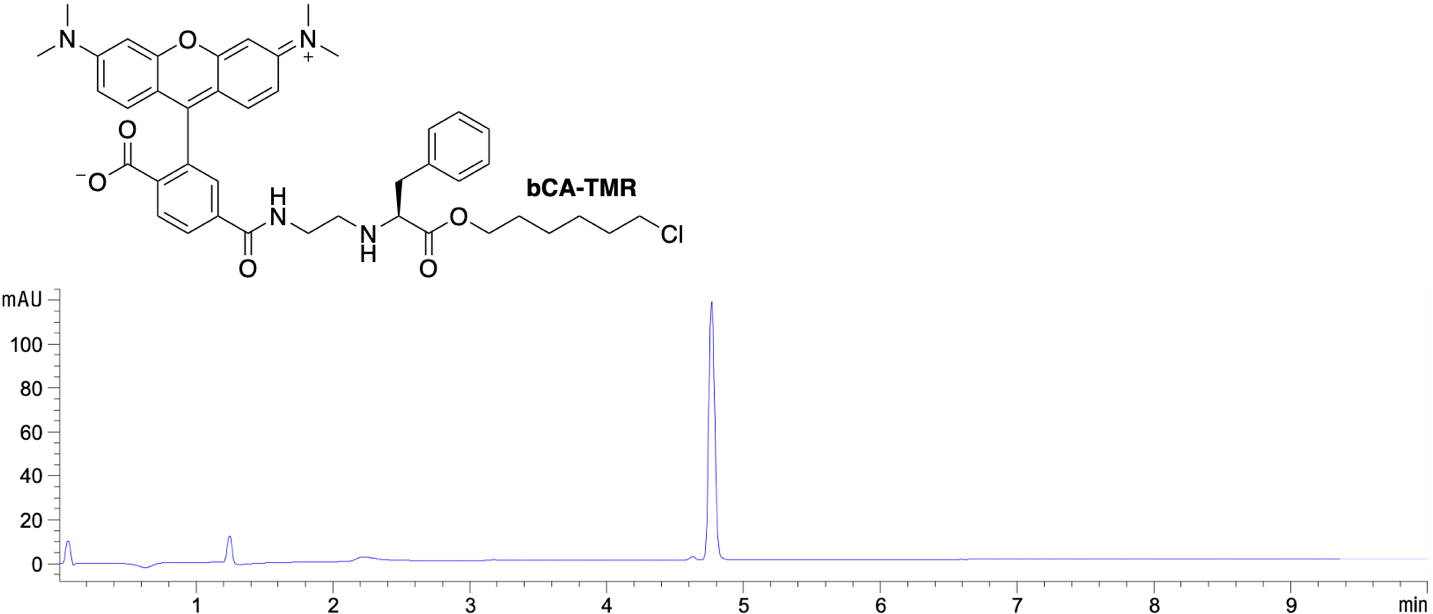

HPLC characterization of bCA-TMR at 545 nm. LC-MS (ESI+) exact mass calculated for C_42_H_48_ClN_4_O_6_ (MH^+^) 739.32, found 739.3364.

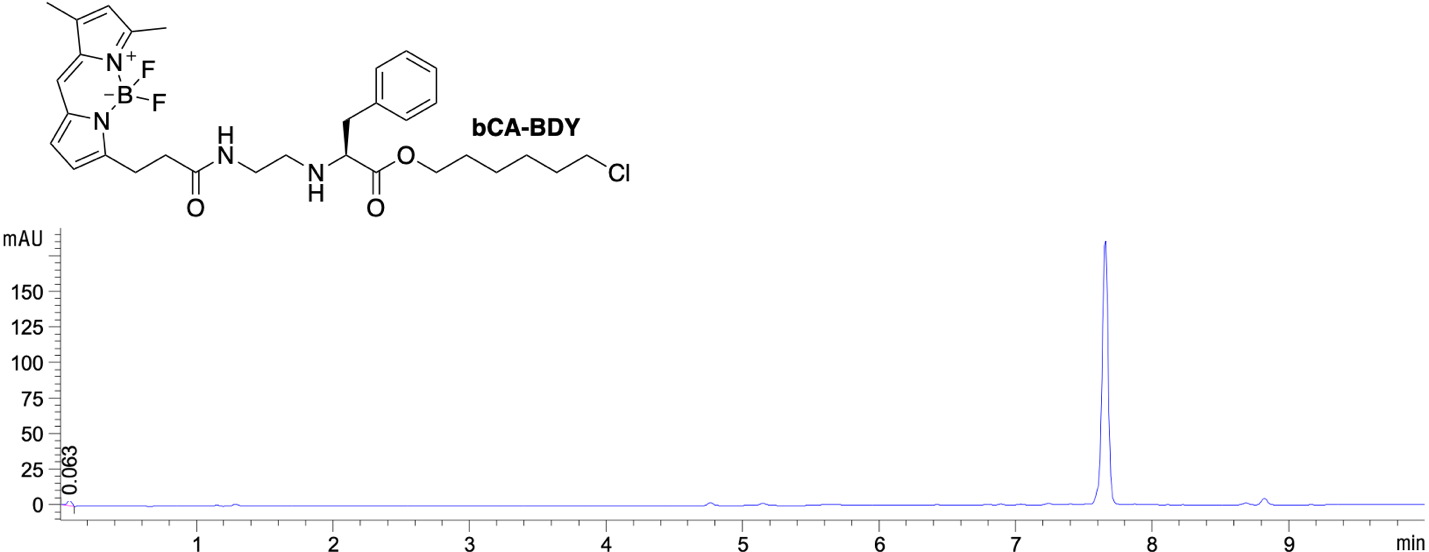

HPLC characterization of bCA-BDY at 495 nm. LC-MS (ESI+) exact mass calculated for C_31_H_41_BClF_2_N_4_O_3_ (MH^+^) 601.29, found 601.3019.

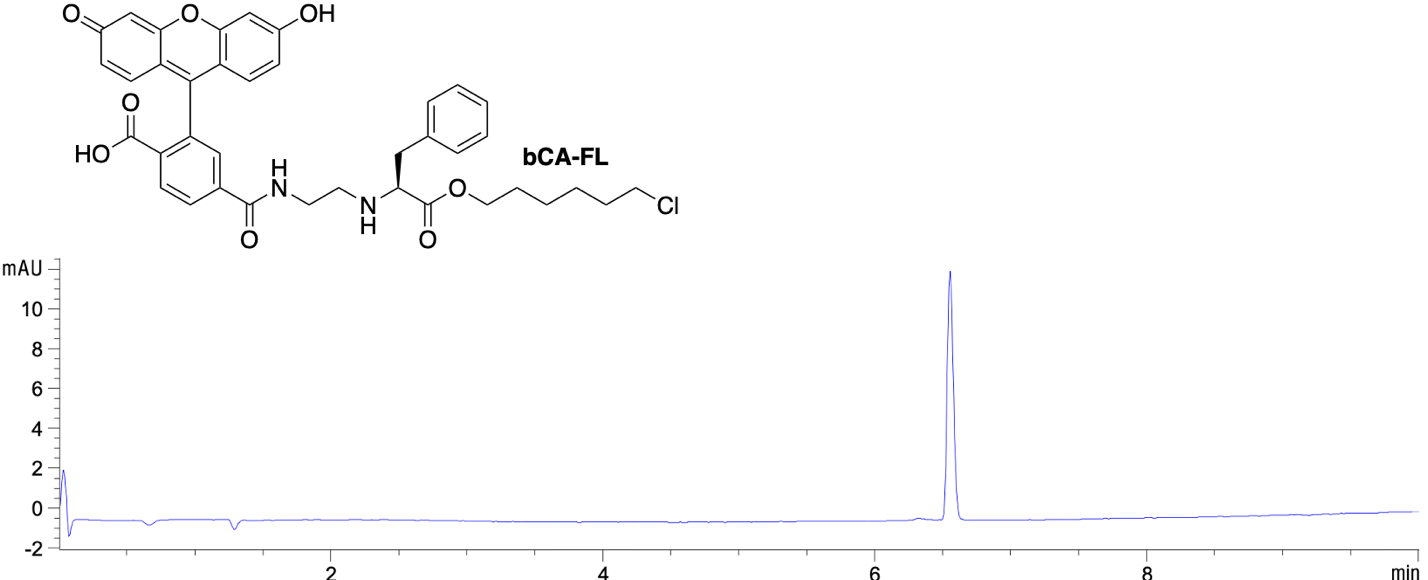

HPLC characterization of bCA-FL at 495 nm. LC-MS (ESI+) exact mass calculated for C_38_H_38_ClN_2_O_8_ (MH^+^) 685.22, found 685.2339.

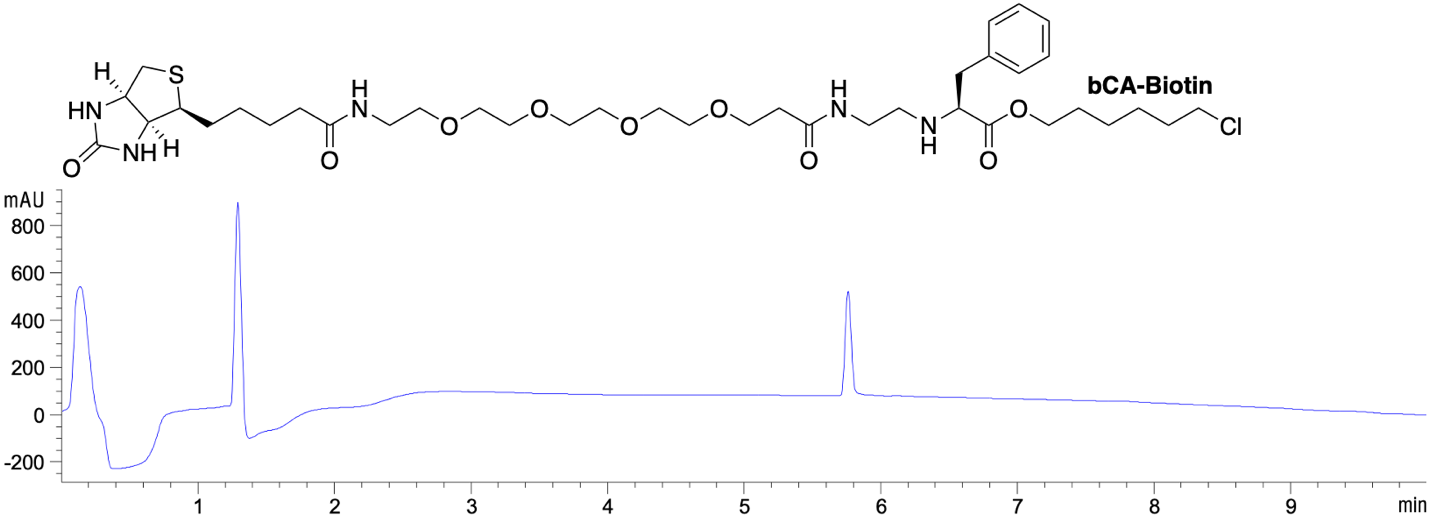

HPLC characterization of bCA-Biotin at 214 nm. LC-MS (ESI+) exact mass calculated for C_38_H_63_ClN_5_O_9_S (MH^+^) 800.40, found 800.4290.

**Supplemental Figure 4.** **HPLC traces of purified, dye-conjugated substrates.**

### Bump and Hole Design

#### **Supplemental Table 1. DNA sequences of four HaloTag7 Phe-to-Ala point mutants.** Homologous overhangs to pCTcon2 are shown in orange. Genes were synthesized by Twist Biosciences.

| **Mutant** | **Sequence** |
| --- | --- |
| F144A | GGAGGCGGTAGCGGAGGCGGAGGGTCGGCTAGCTGCGGTGGCGGCGGTATGGCTGAAATTGGTACAGGTTTTCCATTTGATCCACATTACGTTGAAGTTTTGGGTGAAAGAATGCATTACGTTGATGTTGGTCCAAGAGATGGTACACCAGTTTTGTTTTTACATGGTAACCCAACTTCTTCATACGTTTGGAGAAACATCATCCCACATGTTGCACCAACTCATAGATGTATTGCTCCAGATTTGATTGGTATGGGTAAATCTGATAAGCCAGATTTGGGTTATTTCTTTGATGATCATGTTAGATTCATGGATGCTTTTATTGAAGCATTGGGTTTGGAAGAAGTTGTTTTGGTTATTCATGATTGGGGTTCTGCATTAGGTTTTCATTGGGCTAAGAGAAACCCAGAAAGAGTTAAGGGTATCGCTTTTATGGAGTTTATTAGACCAATTCCAACATGGGATGAATGGCCAGAAGCTGCAAGAGAAACTTTCCAAGCTTTTAGAACAACTGATGTTGGTAGAAAGTTGATCATCGATCAAAACGTTTTTATTGAAGGTACATTGCCAATGGGTGTTGTTAGACCATTGACTGAAGTTGAAATGGATCATTACAGAGAACCATTTTTAAACCCAGTTGATAGAGAACCATTGTGGAGATTTCCAAATGAATTACCAATTGCTGGTGAACCAGCAAACATCGTTGCTTTGGTTGAAGAATACATGGATTGGTTACATCAATCTCCAGTTCCAAAGTTGTTATTTTGGGGTACACCAGGTGTTTTAATTCCACCAGCAGAAGCTGCAAGATTGGCTAAGTCATTGCCAAACTGTAAGGCAGTTGATATTGGTCCAGGTTTGAATTTGTTGCAAGAAGATAACCCAGATTTGATTGGTTCTGAAATTGCTAGATGGTTGTCAACTTTAGAAATTTCTGGGGGCGGATCCGAACAAAAGCTTATTTCTGAAGAGGAC |
| F149A | GGAGGCGGTAGCGGAGGCGGAGGGTCGGCTAGCTGCGGTGGCGGCGGTATGGCTGAAATTGGTACAGGTTTTCCATTTGATCCACATTACGTTGAAGTTTTGGGTGAAAGAATGCATTACGTTGATGTTGGTCCAAGAGATGGTACACCAGTTTTGTTTTTACATGGTAACCCAACTTCTTCATACGTTTGGAGAAACATCATCCCACATGTTGCACCAACTCATAGATGTATTGCTCCAGATTTGATTGGTATGGGTAAATCTGATAAGCCAGATTTGGGTTATTTCTTTGATGATCATGTTAGATTCATGGATGCTTTTATTGAAGCATTGGGTTTGGAAGAAGTTGTTTTGGTTATTCATGATTGGGGTTCTGCATTAGGTTTTCATTGGGCTAAGAGAAACCCAGAAAGAGTTAAGGGTATCGCTTTTATGGAGTTTATTAGACCAATTCCAACATGGGATGAATGGCCAGAATTTGCAAGAGAAACTGCTCAAGCTTTTAGAACAACTGATGTTGGTAGAAAGTTGATCATCGATCAAAACGTTTTTATTGAAGGTACATTGCCAATGGGTGTTGTTAGACCATTGACTGAAGTTGAAATGGATCATTACAGAGAACCATTTTTAAACCCAGTTGATAGAGAACCATTGTGGAGATTTCCAAATGAATTACCAATTGCTGGTGAACCAGCAAACATCGTTGCTTTGGTTGAAGAATACATGGATTGGTTACATCAATCTCCAGTTCCAAAGTTGTTATTTTGGGGTACACCAGGTGTTTTAATTCCACCAGCAGAAGCTGCAAGATTGGCTAAGTCATTGCCAAACTGTAAGGCAGTTGATATTGGTCCAGGTTTGAATTTGTTGCAAGAAGATAACCCAGATTTGATTGGTTCTGAAATTGCTAGATGGTTGTCAACTTTAGAAATTTCTGGGGGCGGATCCGAACAAAAGCTTATTTCTGAAGAGGAC |
| F152A | GGAGGCGGTAGCGGAGGCGGAGGGTCGGCTAGCTGCGGTGGCGGCGGTATGGCTGAAATTGGTACAGGTTTTCCATTTGATCCACATTACGTTGAAGTTTTGGGTGAAAGAATGCATTACGTTGATGTTGGTCCAAGAGATGGTACACCAGTTTTGTTTTTACATGGTAACCCAACTTCTTCATACGTTTGGAGAAACATCATCCCACATGTTGCACCAACTCATAGATGTATTGCTCCAGATTTGATTGGTATGGGTAAATCTGATAAGCCAGATTTGGGTTATTTCTTTGATGATCATGTTAGATTCATGGATGCTTTTATTGAAGCATTGGGTTTGGAAGAAGTTGTTTTGGTTATTCATGATTGGGGTTCTGCATTAGGTTTTCATTGGGCTAAGAGAAACCCAGAAAGAGTTAAGGGTATCGCTTTTATGGAGTTTATTAGACCAATTCCAACATGGGATGAATGGCCAGAATTTGCAAGAGAAACTTTTCAAGCTGCTAGAACAACTGATGTTGGTAGAAAGTTGATCATCGATCAAAACGTTTTTATTGAAGGTACATTGCCAATGGGTGTTGTTAGACCATTGACTGAAGTTGAAATGGATCATTACAGAGAACCATTTTTAAACCCAGTTGATAGAGAACCATTGTGGAGATTTCCAAATGAATTACCAATTGCTGGTGAACCAGCAAACATCGTTGCTTTGGTTGAAGAATACATGGATTGGTTACATCAATCTCCAGTTCCAAAGTTGTTATTTTGGGGTACACCAGGTGTTTTAATTCCACCAGCAGAAGCTGCAAGATTGGCTAAGTCATTGCCAAACTGTAAGGCAGTTGATATTGGTCCAGGTTTGAATTTGTTGCAAGAAGATAACCCAGATTTGATTGGTTCTGAAATTGCTAGATGGTTGTCAACTTTAGAAATTTCTGGGGGCGGATCCGAACAAAAGCTTATTTCTGAAGAGGAC |
| F168A | GGAGGCGGTAGCGGAGGCGGAGGGTCGGCTAGCTGCGGTGGCGGCGGTATGGCTGAAATTGGTACAGGTTTTCCATTTGATCCACATTACGTTGAAGTTTTGGGTGAAAGAATGCATTACGTTGATGTTGGTCCAAGAGATGGTACACCAGTTTTGTTTTTACATGGTAACCCAACTTCTTCATACGTTTGGAGAAACATCATCCCACATGTTGCACCAACTCATAGATGTATTGCTCCAGATTTGATTGGTATGGGTAAATCTGATAAGCCAGATTTGGGTTATTTCTTTGATGATCATGTTAGATTCATGGATGCTTTTATTGAAGCATTGGGTTTGGAAGAAGTTGTTTTGGTTATTCATGATTGGGGTTCTGCATTAGGTTTTCATTGGGCTAAGAGAAACCCAGAAAGAGTTAAGGGTATCGCTTTTATGGAGTTTATTAGACCAATTCCAACATGGGATGAATGGCCAGAATTTGCAAGAGAAACTTTTCAAGCTTTTAGAACAACTGATGTTGGTAGAAAGTTGATCATCGATCAAAACGTTGCTATTGAAGGTACATTGCCAATGGGTGTTGTTAGACCATTGACTGAAGTTGAAATGGATCATTACAGAGAACCATTTTTAAACCCAGTTGATAGAGAACCATTGTGGAGATTTCCAAATGAATTACCAATTGCTGGTGAACCAGCAAACATCGTTGCTTTGGTTGAAGAATACATGGATTGGTTACATCAATCTCCAGTTCCAAAGTTGTTATTTTGGGGTACACCAGGTGTTTTAATTCCACCAGCAGAAGCTGCAAGATTGGCTAAGTCATTGCCAAACTGTAAGGCAGTTGATATTGGTCCAGGTTTGAATTTGTTGCAAGAAGATAACCCAGATTTGATTGGTTCTGAAATTGCTAGATGGTTGTCAACTTTAGAAATTTCTGGGGGCGGATCCGAACAAAAGCTTATTTCTGAAGAGGAC |

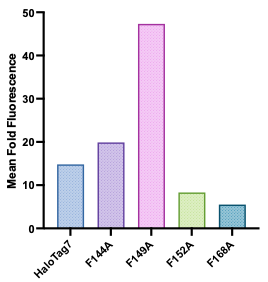

#### **Supplemental Figure 5. Characterization of four Phe-to-Ala point mutants of HaloTag7 using yeast display.** Mean fold fluorescence over yeast cell autofluorescence of populations of 5000 yeast cells expressing HaloTag7 or one of four point mutants when labeled with 1 μM bumped chloroalkane BODIPY-FL for 15 minutes, as measured using flow cytometry.

### Protein Expression and Purification

HaloTag7 and HaloTag7 mutants were recombinantly expressed in BL21(DE3) competent *E. coli* (NEB) and purified by batch affinity chromatography on HisPur™️ Ni-NTA resin (Thermo Fisher Scientific) according to prior published protocols.^3^ Protein yields were typically 20-25 mg per liter culture for yeast codon-optimized constructs and up to 90 mg/L for bacterial codon-optimized constructs.

### Enzyme Kinetics of Rationally Designed Variants

Enzyme kinetics were measured using growth of fluorescence polarization over time for 250 nM fluorescent chloroalkane substrate binding to multiple concentrations of excess protein. Data were fit using a pseudo first-order kinetic equation to determine an apparent second order rate constant as previously described.^3^

Equation: $Y=Y_{0}+(Plateau- Y_{0})(1- e^{{-k}_{obs}X})$

#### **Supplemental Table 2. Apparent second-order rate constants of HaloTag7 and F149A reacting with TMR, FL, and BDY substrates.** *HaloTag rate constant with CA-TMR was too fast to be measured using this methodology. We report a value from the literature.^4^

| Protein | Replicate | CA-FL (M^-1^s^-1^) | bCA-FL (M^-1^s^-1^) | CA-TMR (M^-1^s^-1^) | bCA-TMR  (M^-1^s^-1^) | CA-BDY (M^-1^s^-1^) | bCA-BDY (M^-1^s^-1^) |
| --- | --- | --- | --- | --- | --- | --- | --- |
| HaloTag7 | 1 | 51900 | 6.44 | 1.88 x 10^7*^ | 134 | 4490 | 9.1 |
|  | 2 | 42600 | 8.68 | X | 156 | 3990 | 10.2 |
|  | 3 | 50500 | 14.0 | X | 124 | 4800 | 10.6 |
|  | 4 | 27400 | 42.3 | X | 145 | 2330 | 123 |
|  | 5 | 34200 | 112 | X | 115 | 2480 | 127 |
|  | 6 | 48600 | 33.6 | X | 126 | 1390 | 82.1 |
|  | 7 | X | 113 | X | X | X | X |
| F149A | 1 | 236 | 371 | 6560 | 562 | 52.0 | 250 |
|  | 2 | 229 | 356 | 6030 | 727 | 75.4 | 336 |
|  | 3 | 252 | 314 | 7180 | 623 | 39.8 | 248 |
|  | 4 | 329 | 157 | X | 1540 | X | X |
|  | 5 | 326 | 155 | X | 1500 | X | X |
|  | 6 | 412 | 161 | X | X | X |  |

### Library Generation and Directed Evolution

#### **Supplemental Table 3. HaloTag F149A mutagenic sub libraries.**

| **Sub Library** | **Error-Prone PCR Conditions** | **Estimated diversity**  **(x 10^6^)** | **Percent activity relative to F149A** | **Average number of amino acid mutations per gene** |
| --- | --- | --- | --- | --- |
| 1 | 2 μM 8-Oxo-dGTP*^a^*  2 μM dPTP*^b^*  10 cycles | 3.55 | 67% | 1.63 |
| 2 | 10 μM 8-Oxo-dGTP  10 μM dPTP  10 cycles | 6.45 | 19% | 5.94 |
| 3 | 2 μM 8-Oxo-dGTP  2 μM dPTP  20 cycles | 5.95 | 65% | 1.81 |
| 4 | 10 μM 8-Oxo-dGTP  10 μM dPTP  20 cycles | 1.15 | 29% | 8.17 |
| Pooled input library |  | 17.1 | 46% | 3.76 |

*^a^ 8-Oxo-dGTP: 8-oxo-2’-deoxyguanosine-5’-triphosphate; ^b^ dPTP: 2’-deoxy-P-nucleoside-5’-triphosphate*

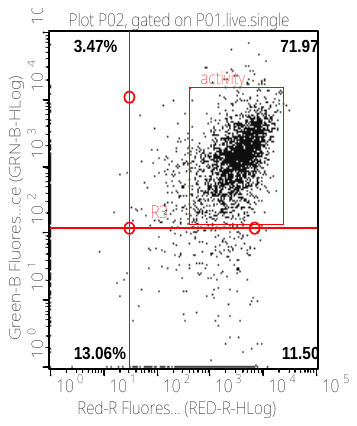

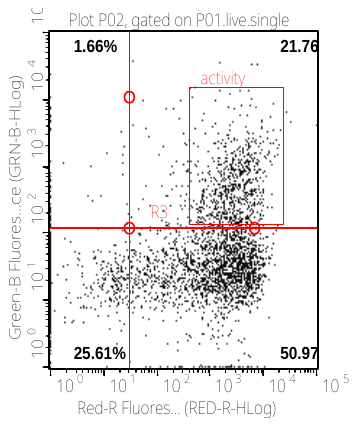

*Sub-library 1	Sub-library 2*

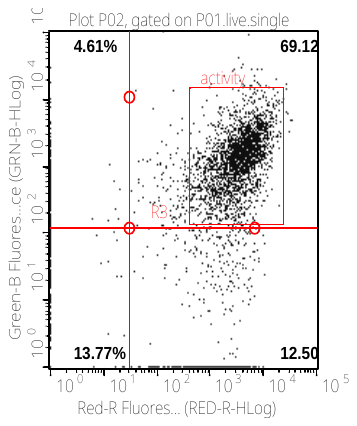

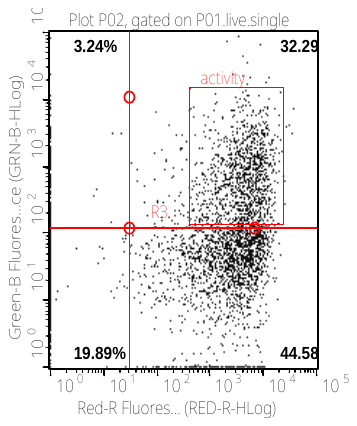

*Sub-library 3	Sub-library 4*

#### **Supplemental Figure 6. Activity of four sub-libraries derived from F149A.** Mutagenized sub-libraries of HaloTag F149A (Table S4) were expressed on the surface of yeast and labeled with 1 μM bumped chloroalkane-BODIPY-FL for 15 minutes before being stained with anti-Myc AF647 antibodies and measured on a flow cytometer. The y-axis shows HaloTag activity through chloroalkane substrate labeling and the x-axis shows expression of the full-length construct by immunostaining against a C-terminal Myc tag. Each dot represents an individual yeast cell and 5000 cells are shown on each plot. The red rectangular gate labeled “activity” is drawn based on the observed labeling of a population of yeast expressing HaloTag F149A under the same conditions.

#### **Supplemental Table 4. Labeling conditions during three rounds of iterative fluorescence-activated cell sorting enrichment to evolve for an orthogonal HaloTag variant.**

| **Round** | **[bCA-FL] (nM)** | **[bCA-biotin] (nM)** | **[CA] (fold excess)** | **Labeling time (min)** |
| --- | --- | --- | --- | --- |
| Input: pooled library of 17.1 million F149A mutants | | | | |
| 1 | 300 | 0 | 3x | 15 |
| 2.1 | 0 | 200 | 30x | 15 |
| 2.2 | 0 | 200 | 300x | 15 |
| 1′ | 0 | 1000 | 3x | 15 |
| 2.1′ | 60 | 0 | 30x | 15 |
| 2.2′ | 60 | 0 | 300x | 15 |
| Pool round 2 populations | | | | |
| 3.1 | 6 | 0 | 300x | 10 |
| 3.2 | 6 | 0 | 300x | 1 |
| 3.1′ | 0 | 20 | 300x | 10 |
| 3.2′ | 0 | 20 | 300x | 1 |
| Output: four populations of ~ 6 thousand highly active mutants | | | | |

**Selection strategy and conditions**

Because each of the mutagenic sub-libraries from our error-prone PCR contained at least 21% active variants (Fig. S6), we pooled them into a single input library with 1.71 x 10^7^ members to use for directed evolution. We performed parallel sorts by labeling with either bCA-FL and then bCA-biotin (rounds 1, 2.1, and 2.2), or bCA-biotin and then bCA-FL (rounds 1′, 2.1′, and 2.2′) (Table S5). An excess of linear chloroalkane with no conjugated dye was applied in every round to enforce selectivity for bumped substrates. Stringency was increased in each successive round by decreasing concentrations of bumped substrates, decreasing labeling times, and increasing concentrations of non-fluorescent linear chloroalkane. At a library size of 17.1 million, we were able to forego physical isolation methods and use fluorescence-activated cell sorting even in the first round. We sorted enough yeast cells to cover at least ten-fold the diversity of the input for each round, except for the first round where ~1 x 10^8^ cells were sorted due to time constraints. The top ~0.05% of brightest cells were isolated in the first round, and the top ~0.5-1.0% of cells were isolated in subsequent rounds. Rounds 2.1/2.2 and 2.1′/2.2′ were performed in parallel with different stringencies. Because all round 2 output populations showed a similar increase in activity, they were pooled into a single population before round 3. Four different labeling conditions were applied in round 3, and all round 3 output pools showed robust labeling with bumped substrates in the presence of excess linear chloroalkane (Fig. S9 and Fig. 2b). Thus, we sequenced all round 3 output pools to identify enriched mutations.

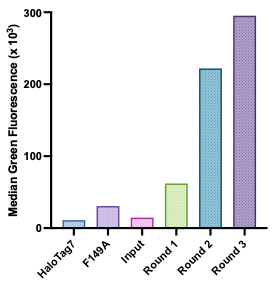

#### **Supplemental Figure 7. Increase in activity of F149A libraries between rounds of enrichment.** The median green fluorescence of HaloTag7, F149A, the pooled input library, and the pooled output populations of three rounds of FACS enrichment was measured using flow cytometry after labeling yeast cells expressing the HaloTag variants with 60 nM bumped chloroalkane-fluorescein and 300-fold excess of nonfluorescent linear chloroalkane for 15 minutes.

#### **Supplemental Table 5. Sequencing of input and output libraries.** DNA was isolated from yeast populations, and five thousand sequences were generated for each library using Plasmidsaurus premium PCR sequencing. Mutations were counted using Geneious Prime by mapping to a reference F149A sequence and identifying single nucleotide polymorphisms.

| Population | Mutation | Frequency |
| --- | --- | --- |
| Library 0 | No mutations > 5% detected | |
| Round 3.1 | P142S | 6.9% |
|  | L173V | 5.3% |
|  | I218T | 5.7% |
|  | N272Y | 99.8% |
| Round 3.1 Prime | T58A | 6.4% |
|  | F91L | 6.9% |
|  | G96S | 7% |
|  | P142S | 14.9% |
|  | F144L | 9% |
|  | F144S | 7.1% |
|  | D187V | 5.1% |
|  | I218T | 11% |
|  | N272Y | 99% |
| Round 3.2 | L173V | 5.3% |
|  | N272Y | 96.5% |
| Round 3.2 Prime | F91L | 7.1% |
|  | G96S | 6.7% |
|  | P142S | 27% |
|  | F144L | 12.2% |
|  | D187V | 10.1% |
|  | I218T | 18.2% |
|  | N272Y | 93.7% |

#### **Supplemental Table 6. Co-occurring mutations in output libraries.** To look for co-occurring mutations, sequence groups containing the most common mutations (P142S and F144L) were created for each output library and examined for single nucleotide polymorphisms (SNP) using Geneious Prime after mapping to a reference F149A sequence. Mutations which co-occurred with each other within sequence groups are color coded, excluding N272Y which occurred in virtually all sequences.

| 3.1 | | | |
| --- | --- | --- | --- |
| Sequence Group | SNP | AA Mutation | Frequency |
| P142S | A159T | Silent | 23.2% |
|  | A212G | K71R | 23.4% |
|  | T303C | Silent | 68.4% |
|  | T552C | Silent | 66.4% |
|  | A560T | D187V | 25.8% |
|  | T653C | I218T | 64.3% |
|  | A753G | Silent | 21.9% |
|  | A814T | N272Y | 100.0% |
|  | T852A | Silent | 11.2% |

| 3.1 Prime | | | |
| --- | --- | --- | --- |
| Sequence Group | SNP | AA Mutation | Frequency |
| P142S | A159T | Silent | 26.9% |
|  | A212G | K71R | 28.2% |
|  | T303C | Silent | 61.4% |
|  | T552C | Silent | 61.0% |
|  | A560T | D187V | 32.6% |
|  | T653C | I218T | 60.0% |
|  | A753G | Silent | 29.5% |
|  | A814T | N272Y | 99.4% |
|  | T852A | Silent | 7.7% |
| F144L | T45A | Silent | 15.9% |
|  | A48G | Silent | 16.5% |
|  | TT270-271CC | F91L | 70.6% |
|  | G286A | G96S | 70.1% |
|  | A814T | N272Y | 97.9% |
|  | T852A | Silent | 23.0% |

| 3.2 Prime | | | |
| --- | --- | --- | --- |
| Sequence Group | SNP | AA Mutation | Frequency |
| P142S | A159T | Silent | 33.9% |
|  | A212G | K71R | 34.3% |
|  | T303C | Silent | 57.1% |
|  | T552C | Silent | 57.3% |
|  | A560T | D187V | 36.2% |
|  | T653C | I218T | 57.4% |
|  | A753G | Silent | 34.6% |
|  | A814T | N272Y | 98.7% |
|  | T852A | Silent | 7.8% |
| F144L | A36G | Silent | 6.5% |
|  | T45A | Silent | 19.4% |
|  | A48G | Silent | 18.0% |
|  | T255C | Silent | 5.9% |
|  | TT270-271CC | F91L | 51.3% |
|  | G286A | G96S | 51.1% |
|  | A514G | T172A | 13.0% |
|  | G529A | V177I | 6.0% |
|  | A705G | Silent | 5.7% |
|  | A753G | Silent | 6.9% |
|  | A814T | N272Y | 84.7% |
|  | T852A | Silent | 28.4% |

### Enzyme Kinetics of Evolved Variants

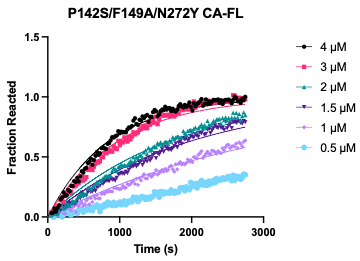

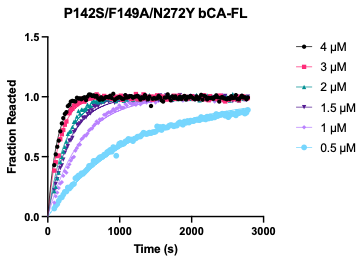

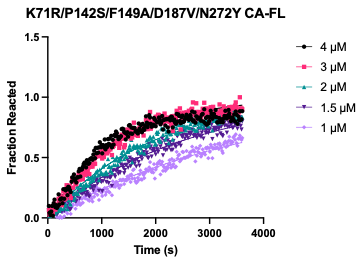

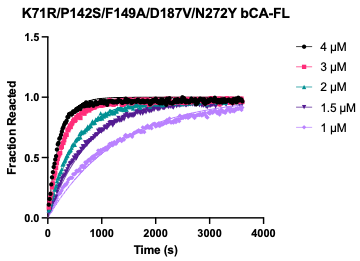

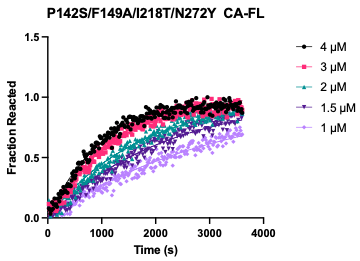

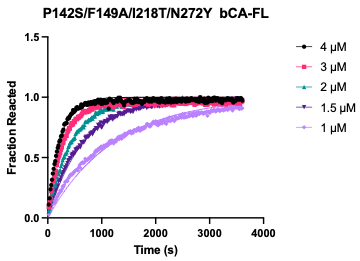

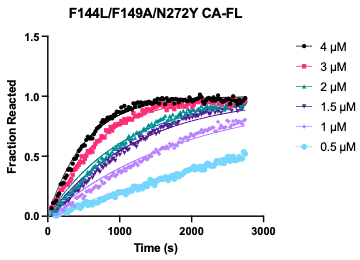

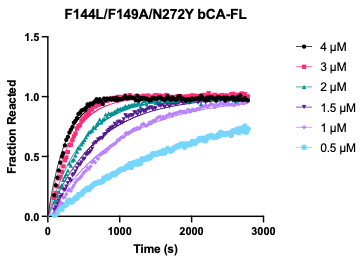

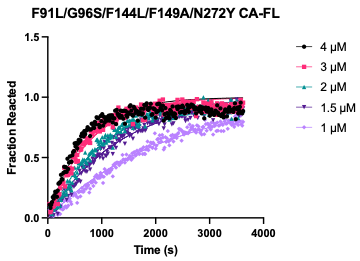

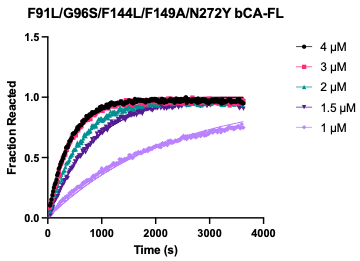

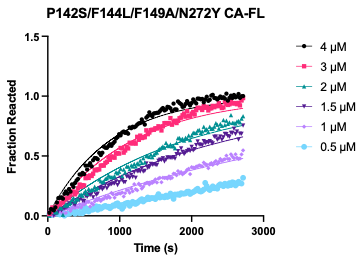

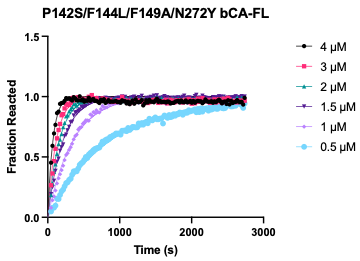

#### **Supplemental Figure 8.** **Selected representative, normalized fluorescence polarization kinetic traces of HaloTag mutants with fluorescein-conjugated substrates.** One of three total replicates with one-phase association curve fits are shown. Data are normalized to the saturation value for each mutant/substrate pair.

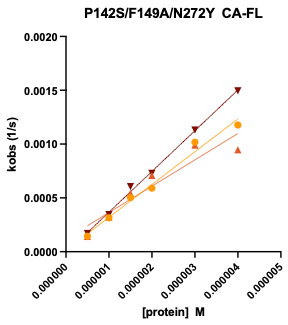

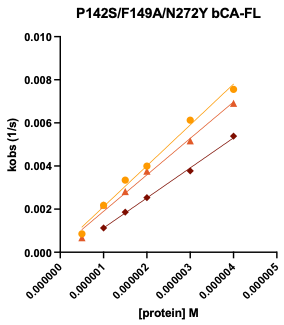

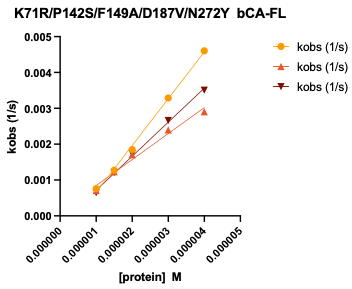

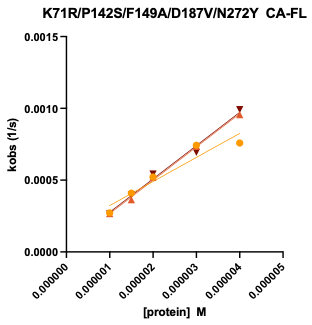

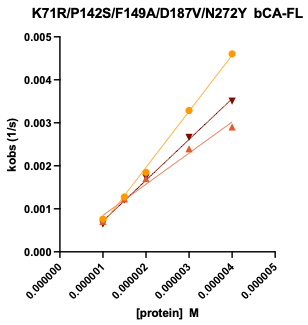

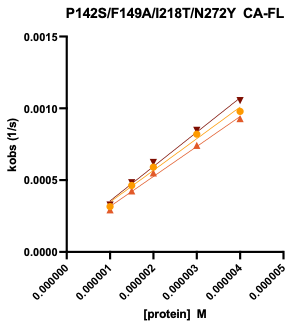

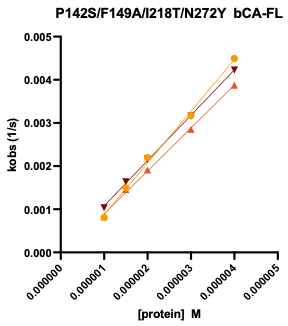

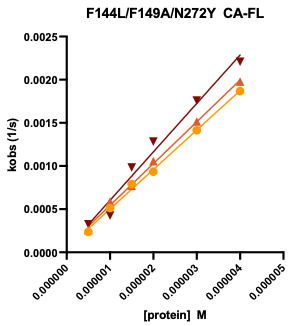

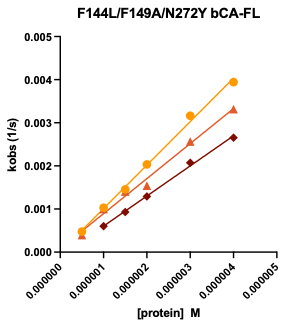

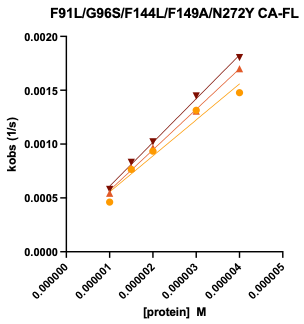

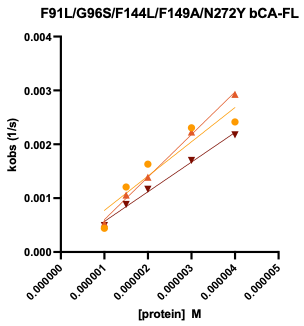

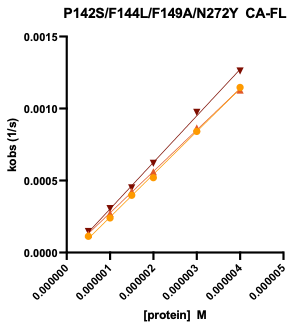

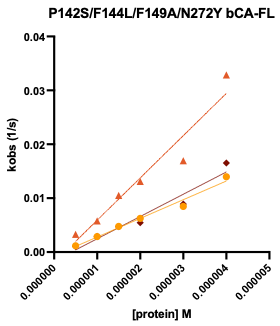

#### **Supplemental Figure 9. Selected 2^nd^ order rate constant plots.** Observed rate constants are plotted against protein concentration to generate apparent second-order rate constants from the slope of the fitted line.

#### **Supplemental Table 7.** **Apparent second-order rate constants of HaloTag7 and OrthoTag with TMR and BDY substrates.** Data are shown as the average of three replicates with the standard error of the mean.

| Protein | k_app_ CA-TMR (M^-1^s^-1^) | k_app_ bCA-TMR (M^-1^s^-1^) | k_app_ CA-BDY (M^-1^s^-1^) | k_app_ bCA-BDY (M^-1^s^-1^) |
| --- | --- | --- | --- | --- |
| HaloTag | - | 133 ± 6 | 3250 ± 559 | 60.4 ± 23.5 |
| OrthoTag | 3780 ± 110 | 5310 ± 1240 | 353 ± 76 | 440 ± 58 |

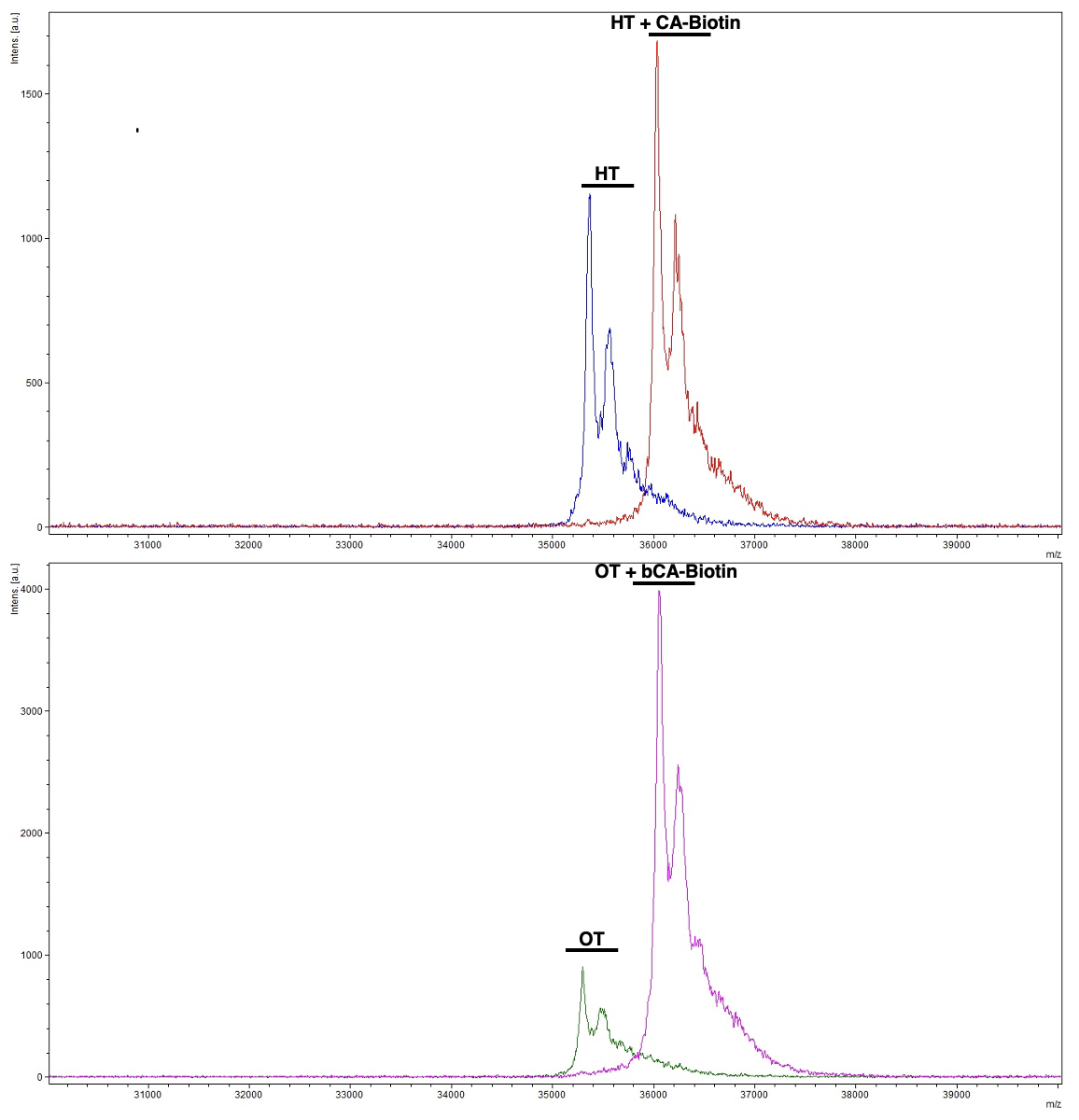

#### **Supplemental Figure 10. Measurement of covalent labeling of purified proteins by mass spectrometry.** 10 µM purified OrthoTag or HaloTag was reacted with 20 µM bCA-biotin or CA-biotin (Promega) respectively at room temperature for 3 h in PBS. Covalent complex formation was observed by MALDI-TOF MS using sinapic acid matrix (Millipore Sigma).

### Crystallography

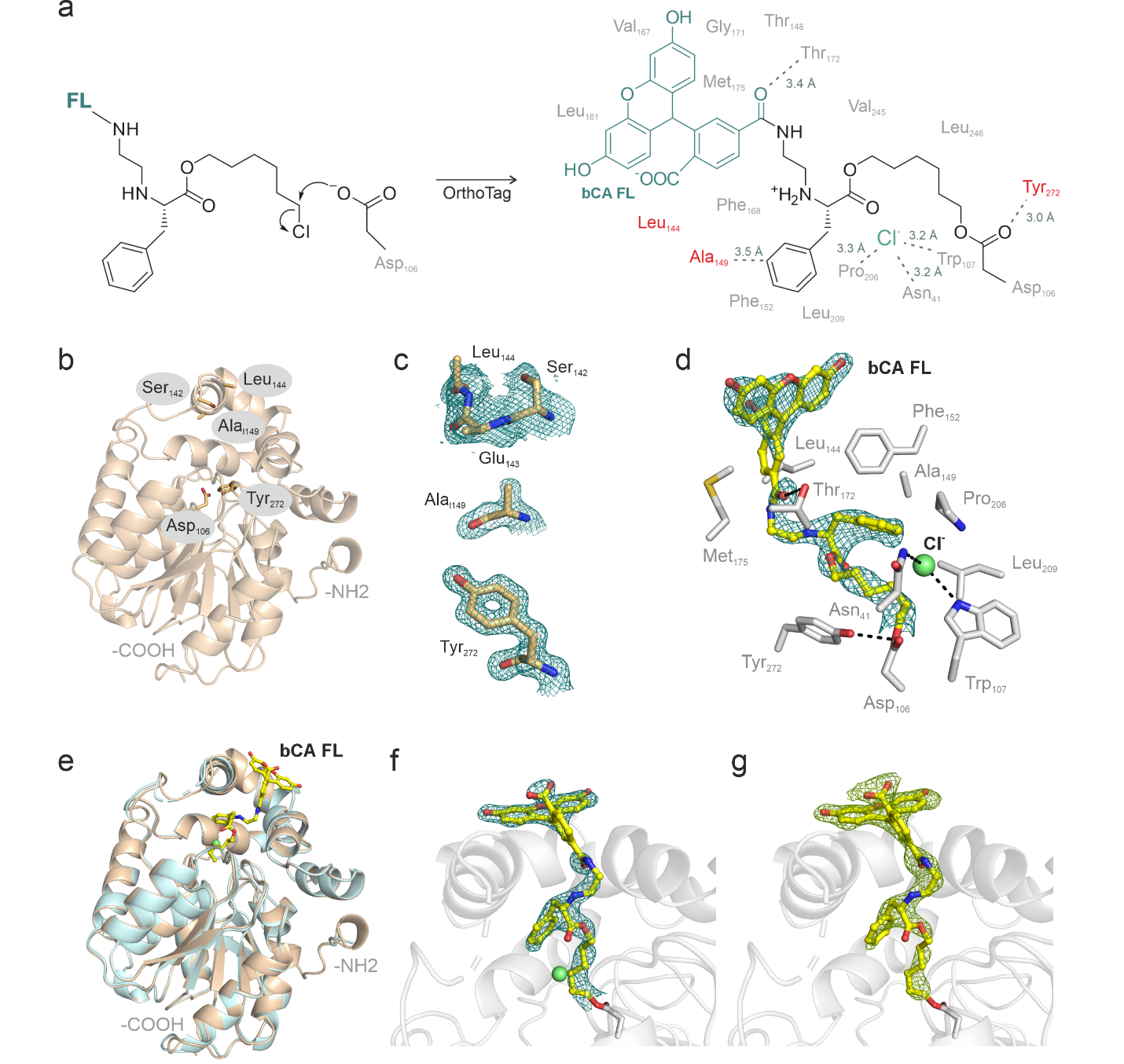

#### **Supplemental Figure 11. OrthoTag crystal structure substrate interactions and electron density.** A) Schematic reaction scheme for covalent bond formation between bCA-FL and OrthoTag. Residues proximal to and interacting with the ligand are shown, with OrthoTag specific mutations labelled red. B) Overview of apo OrthoTag structure (PDB: 30HW) showing position of OrthoTag mutations and the key catalytic residue Asp106. C) 2Fo-Fc maps of OrthoTag derived mutations (PDB: 30HW, 1.40 Å resolution, σ = 1.0) showing electron density surrounding mutated residues. The electron density was poorly resolved for Leu144, therefore the structure was modelled with the sidechain truncated down to the first carbon. D) 2Fo-Fc map (PDB: 30HV, 2.10 Å resolution, σ = 1.0) of bCA-FL with active site proximal and interacting residues. E) Structural alignment between apo OrthoTag (PDB: 30HW) and bCA-FL-bound OrthoTag (PDB: 30HV) revealing an overall nearly identical fold (RMSD=0.293 Å). F – G) 2Fo-Fc (left, 2.10 Å resolution, σ = 1.0) and polder omit (right, 2.10 Å resolution, σ = 3.0) maps of bCA-FL in OrthoTag active site (PDB: 30HV)

#### **Supplemental Table 8. OrthoTag crystallography data collection and refinement statistics.**

| Datasets | Apo OrthoTag  (PDB ID: 30HW) | bCA-FL-bound OrthoTag  (PDB ID: 30HV) |
| --- | --- | --- |
| Data Collection (T in K) | Synchrotron, Cryogenic (100) | Synchrotron, Cryogenic (100) |
| Beamline (Wavelength, Å) | Diamond Light Source I03 (0.9763) | Diamond Light Source I04 (0.9537) |
| Detector | DECTRIS EIGER2 XE 16M | DECTRIS EIGER2 XE 16M |
| Data Processing | Xia2.Dials | Xia2.Dials |
| Space group | *P* 1 2_1_ 1 | P 2 2_1_ 2_1_ |
| Cell dimensions |  |  |
| a,b,c (Å) | 43.479, 69.918, 47.602 | 47.442, 69.310, 99.654 |
| α, β, γ (ᵒ) | 90, 111.68, 90 | 90, 90, 90 |
| No. of molecules/ASU | 1 | 1 |
| No. observed reflections | 359546 (17851)* | 264918 (11659) * |
| No. unique reflections | 51718 (2527)* | 19816 (953)* |
| Resolution (Å) | 69.92-1.40 (1.42-1.40)* | 56.90-2.10 (2.14-2.10)* |
| I/σI | 10.2 (0.7)* | 5.9 (0.6)* |
| CC-half | 0.998 (0.492)* | 0.994 (0.540)* |
| Completeness (%) | 99.3 (97.7)* | 100.0 (100.0)* |
| Multiplicity | 7.0 (7.1)* | 13.4(12.2)* |
| Wilson B value (Å^2^) | 12.320 | 26.62 |
| RMS bonds (Å^2^) | 0.007 | 0.002 |
| RMS angles (°) | 0.95 | 0.55 |
| Refinement | PHENIX | PHENIX |
| R_work_/R_free+_ | 0.1672/0.1850 | 0.2028 /0.2422 |
| No. atoms | 2738 | 2450 |
| - Enzyme | 2441 | 2294 |
| - ligand | n.a. | 48 |
| - Water | 293 | 102 |
| Average B-factors | 22.15 | 48.55 |
| - Enzyme | 20.80 | 47.91 |

*Data in highest resolution shell

### Photophysical Characterization

Relative fluorescence quantum yield measurements were gathered using an Agilent Cary 60 UV/VIS spectrophotometer and Horiba Fluoromax spectrofluorometer following published protocols.^5^ First, a standard sample of Rhodamine 6G (Millipore Sigma) was diluted in 99.5% ACS-grade ethanol (Fisher Scientific) in a UV quartz cuvette (FireFly Scientific) until the maximum absorbance peak was measured at below 0.1. Then, CA-TMR or bCA-TMR substrates were diluted into PBS until the maximum absorbance peak was measured at below 0.1 and the absorbance spectrum overlapped with the standard sample. The total photon emission spectra were then collected for both the rhodamine 6G standard and the TMR substrate sample using the overlapping wavelength as ƛ_ex_. Relative quantum yield was calculated from the integral photon flux using the following equation after normalizing each sample to a solvent blank:

$$\Phi_{f,x}= \Phi_{f,st} \cdot\frac{F_{x}}{F_{st}} \cdot\frac{n_{x}^{2}(ƛ_{em})}{n_{st}^{2}(ƛ_{em})}$$

Where $\Phi_{f,x}$ is the relative quantum yield of the sample, $\Phi_{f,st}$ is the known quantum yield of the standard^6^, $\frac{F_{x}}{F_{st}}$ is the ratio of integral photon flux of the sample and standard, and $\frac{n_{x}^{2}(ƛ_{em})}{n_{st}^{2}(ƛ_{em})}$ is the ratio of the refractive indices of the sample solvent and standard solvent.

HaloTag and OrthoTag bound samples were allowed to react for 30 min at room temperature with shaking in a 5-fold excess of protein relative to substrate prior to absorbance and fluorescence readings, which, based on our fluorescence polarization kinetic measurements, would result in complete reaction of the substrate.

#### **Supplemental Table 9. Relative quantum yields of TMR substrates in solution and bound to HaloTag7 or OrthoTag.** Data are shown as the average of 3 replicates with the standard error of the mean.

| **Sample** | **Protein** | **Solvent** | **Φ_f,x_** |
| --- | --- | --- | --- |
| CA-TMR | None | PBS | 0.287 ± 0.004 |
| CA-TMR | HaloTag7 | PBS | 0.352 ± 0.006 |
| bCA-TMR | None | PBS | 0.262 ± 0.011 |
| bCA-TMR | OrthoTag | PBS | 0.270 ± 0.011 |

### Calculation of Pocket Volumes

Pocket volumes for HaloTag and OrthoTag crystal structures were computed using the POVME3 software^7^ (available as a Python library through PyPI) by mapping inclusion spheres to the atoms in the bound chloroalkane substrate and setting the seed sphere at the catalytic D106 residue. Pocket volumes for substrate-bound structures were calculated by first removing the substrate in PyMOL and then running the software on the stripped structure. For OrthoTag, an exclusion sphere was placed at the mouth of the pocket to prevent inflated volume calculations due to the deep surface channel which passes through the mouth of the active site pocket.

#### **Supplemental Table 10. Effects of alterations to POVME3 parameters.** Pocket volume calculations were repeated with each of the following perturbations. Changes in pocket volume calculations were not large enough to alter overall conclusions.

| Perturbation | Center (x, y, z) | Radius (Å) | Calculated Volume (Å³) | Δ from original |
| --- | --- | --- | --- | --- |
| *Holo* OrthoTag  (Original) | [3.0, −24.0, 19.0] | 4.0 | 310 | — |
| Shift +1 Å (x) | [4.0, −24.0, 19.0] | 4.0 | 304 | −1.9% |
| Shift −1 Å (x) | [2.0, −24.0, 19.0] | 4.0 | 331 | +6.8% |
| Shift +1 Å (y) | [3.0, −23.0, 19.0] | 4.0 | 313 | +1.0% |
| Shift −1 Å (y) | [3.0, −25.0, 19.0] | 4.0 | 332 | +7.1% |
| Shift +1 Å (z) | [3.0, −24.0, 20.0] | 4.0 | 335 | +8.1% |
| Shift −1 Å (z) | [3.0, −24.0, 18.0] | 4.0 | 297 | −4.2% |
| Smaller radius | [3.0, −24.0, 19.0] | 3.0 | 354 | +14.2% |
| Larger radius | [3.0, −24.0, 19.0] | 5.0 | 289 | −6.8% |

| POVME3 parameters used to calculate the HaloTag pocket volume:  grid_spacing: 1.0  points_inclusion_sphere:  center: [5.048, -12.910, 11.971]  radius: 4.0  center: [5.088, -14.399, 12.351]  radius: 4.0  center: [5.394, -14.451, 13.759]  radius: 4.0  center: [5.783, -15.906, 14.125]  radius: 4.0  center: [5.990, -15.731, 15.652]  radius: 4.0  center: [5.783, -18.871, 15.594]  radius: 4.0  center: [6.314, -20.163, 16.365]  radius: 4.0  center: [4.530, -21.549, 17.954]  radius: 4.0  center: [3.056, -21.827, 17.927]  radius: 4.0  center: [2.260, -22.226, 20.321]  radius: 4.0  center: [2.722, -22.669, 19.101]  radius: 4.0  contiguous_pocket_seed_sphere:  center: [5.048, -12.910, 11.971]  radius: 3.0  contiguous_points_criteria: 3  convex_hull_exclusion: true  distance_cutoff: 1.09  save_points: true  save_individual_pocket_volumes: true  save_pocket_volumes_trajectory: true  save_volumetric_density_map: true  output_equal_num_points_per_frame: false  n_cores: 4  compress_output: false | POVME3 parameters used to calculate the OrthoTag pocket volume:  grid_spacing: 1.0  points_inclusion_sphere:  center: [5.048, -12.910, 11.971]  radius: 4.0  center: [5.088, -14.399, 12.351]  radius: 4.0  center: [5.394, -14.451, 13.759]  radius: 4.0  center: [5.783, -15.906, 14.125]  radius: 4.0  center: [5.990, -15.731, 15.652]  radius: 4.0  center: [5.783, -18.871, 15.594]  radius: 4.0  center: [6.314, -20.163, 16.365]  radius: 4.0  center: [4.530, -21.549, 17.954]  radius: 4.0  center: [3.056, -21.827, 17.927]  radius: 4.0  center: [2.219, -19.662, 13.726]  radius: 4.0  center: [1.640, -18.658, 14.487]  radius: 4.0  center: [1.020, -17.578, 13.881]  radius: 4.0  center: [0.969, -17.488, 12.507]  radius: 4.0  points_exclusion_sphere:  center: [3.0, -24.0, 19.0]  radius: 4.0  contiguous_pocket_seed_sphere:  center: [5.048, -12.910, 11.971]  radius: 3.0  contiguous_points_criteria: 3  convex_hull_exclusion: true  distance_cutoff: 1.09  save_points: true  save_individual_pocket_volumes: true  save_pocket_volumes_trajectory: true  save_volumetric_density_map: true  output_equal_num_points_per_frame: false  n_cores: 4  compress_output: false |
| --- | --- |

### Labelling in Mammalian Cells

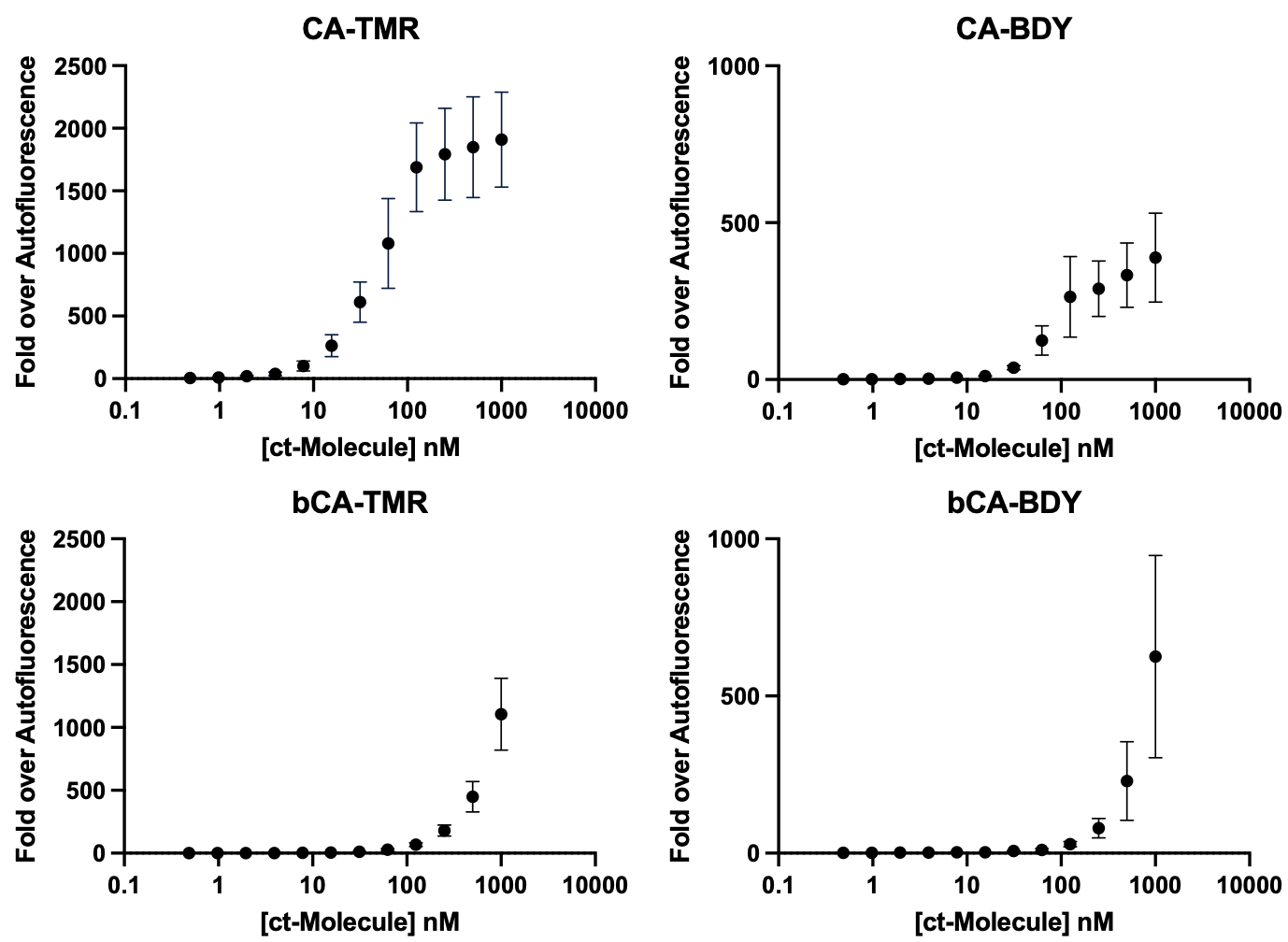

#### **Supplemental Figure 12. Concentration-dependent labeling of HaloTag in mammalian cells with linear and bumped chloroalkane substrates.** HEK293T cells stably expressing HaloTag7 as a fusion to the TOMM20 outer mitochondrial membrane protein were labeled with chloroalkane substrates in OPTI-MEM for 1 hour at 37 °C before washing three times for 15 minutes with OPTI-MEM and measuring the fluorescence of 3000 cells on a flow cytometer. Data points represent the average of three biological replicates and error bars represent the standard error of the mean. Different absolute intensities on different days for background and signal were observed, which is typical for flow cytometry experiments.

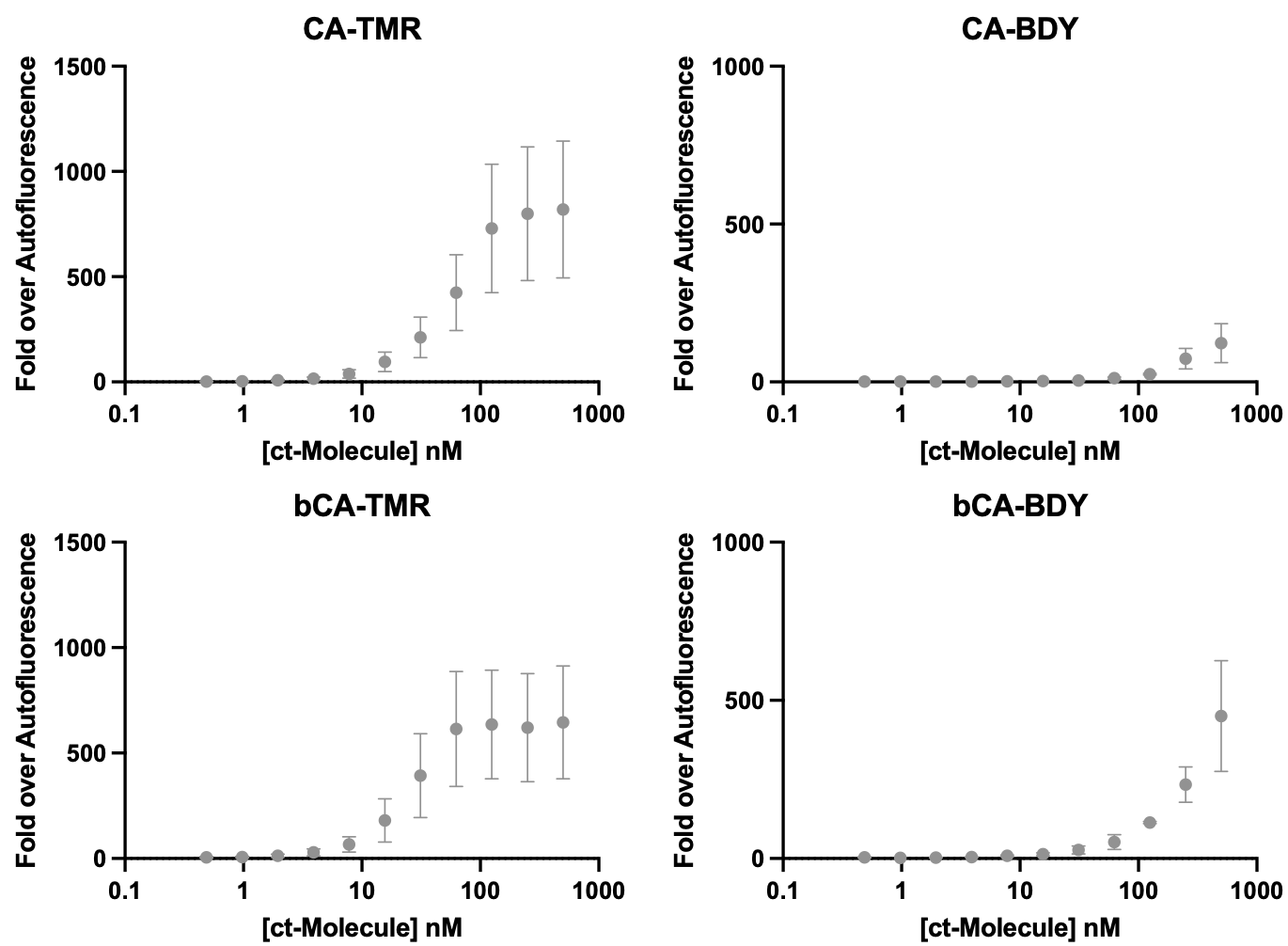

#### **Supplemental Figure 13. Concentration-dependent labeling of OrthoTag in mammalian cells with linear and bumped chloroalkane substrates**. HEK293T cells stably expressing OrthoTag as a fusion to the histone 2B nuclear protein were labeled with chloroalkane substrates in OPTI-MEM for 1 hour at 37 C before washing three times for 15 minutes with OPTI-MEM and measuring the fluorescence of 3000 cells on a flow cytometer. Data points represent the average of three biological replicates and error bars represent the standard error of the mean. Different absolute intensities on different days for background and signal were observed, which is typical for flow cytometry experiments.

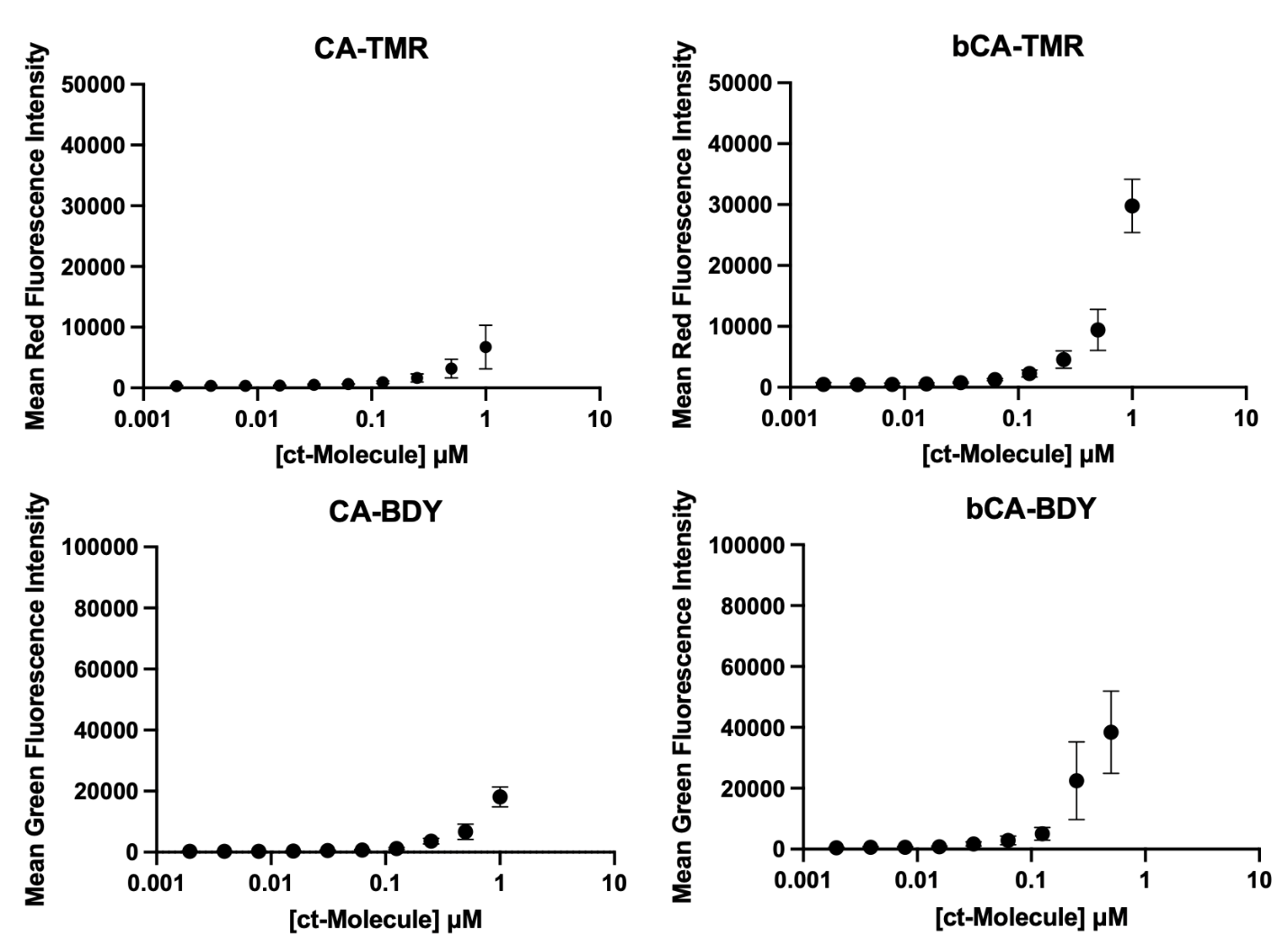

#### **Supplemental Figure 14. Concentration-dependent background signal of CA and bCA TMR and BDY substrates in non-transduced HEK293T cells.** HEK293T cells were labeled with chloroalkane substrates in OPTI-MEM for 1 hour at 37 C before washing three times for 15 minutes with OPTI-MEM and measuring the fluorescence of 3000 cells on a flow cytometer. Data points represent the average of four (TMR substrates) or three (BDY substrates) biological replicates and error bars represent the standard error of the mean. In general, bCA substrates have approximately 2-fold higher background labeling as measured by fluorescence intensity than CA substrates, which we consider a modest increase. Both substrates have minimal background at concentrations of 125 nM and below, which are standard labeling conditions for SLPs in cultured cells. Even at 1 μM concentrations, where the highest background labeling was observed, fluorescence intensities in non-transduced cells are at least 10-fold lower than in cells expressing the corresponding labeling protein.

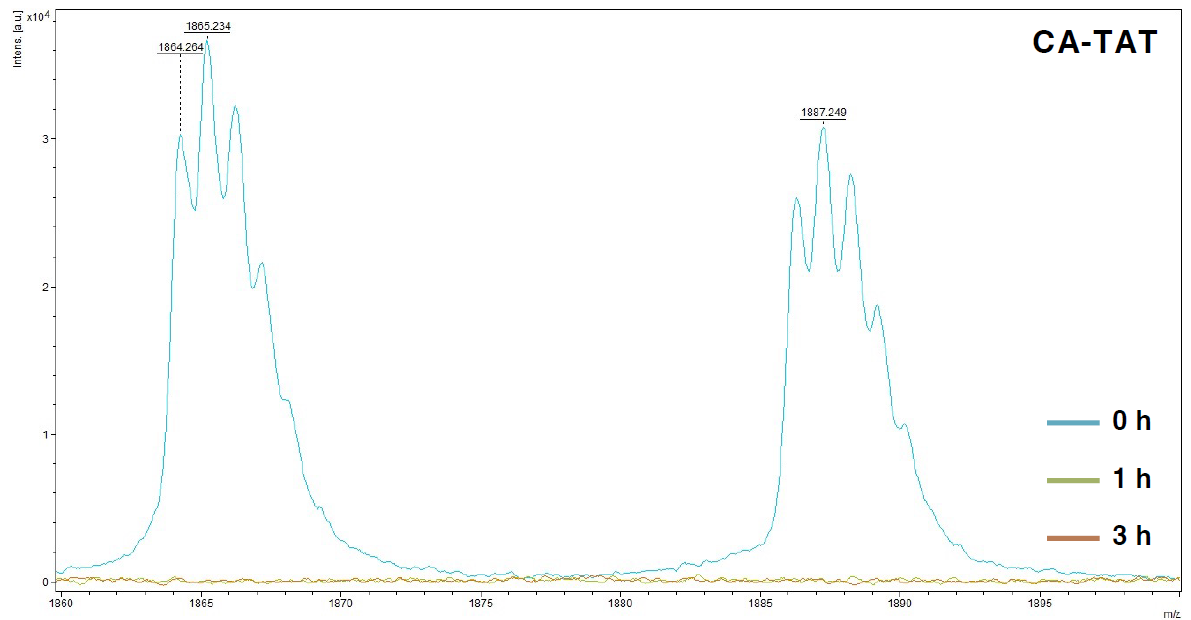

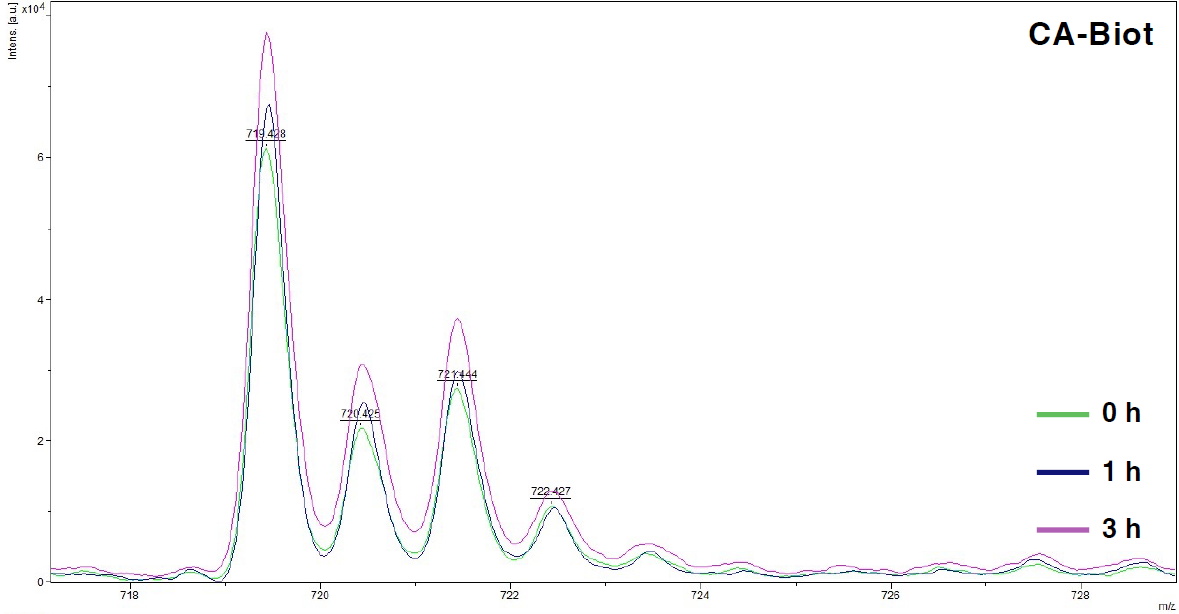

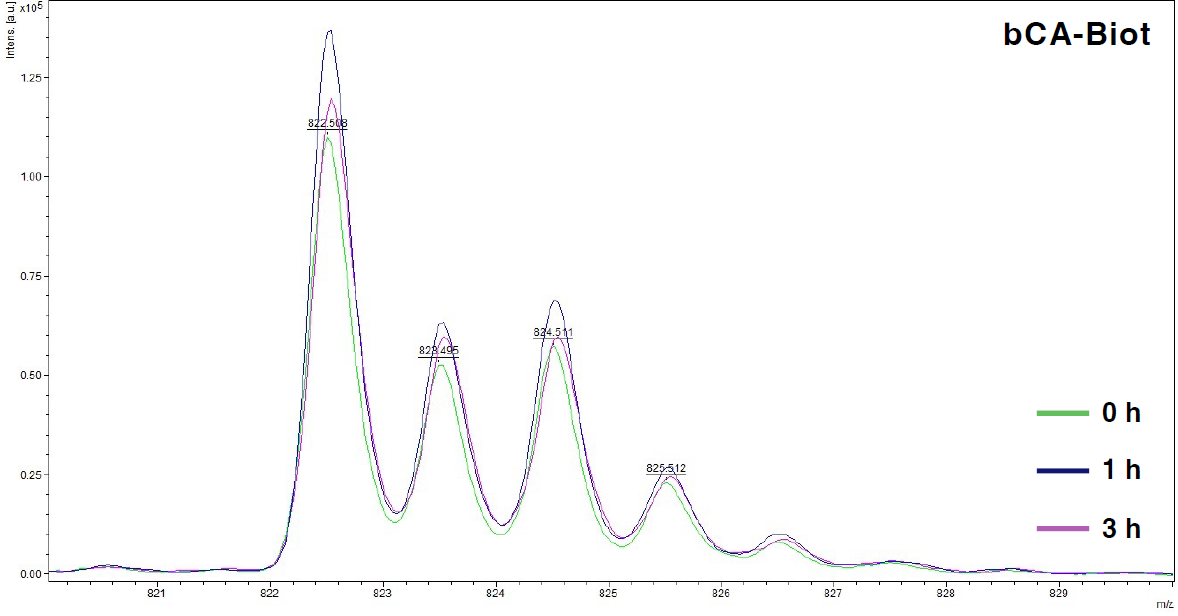

#### **Supplemental Figure 15. MALDI-TOF MS detection of linear and bumped chloroalkane compounds in total cell lysates.** The stability of select compounds in total cell lysates was evaluated according to prior published protocols.^8^ Briefly, 1 x 10^6^ HEK293T cells per compound were washed 2 times in PBS and resuspended in 100 µL lysis buffer (50 mM tris, 250 mM NaCl, 0.5% IGEPAL CA-630, pH 8) and incubated for 15 min on ice before centrifuging for 10 min at 4 ℃ and 21,000 xg and collecting clarified lysate. Compounds were added to lysate to a final concentration of 100 µM in 50 µL while maintaining DMSO less than 0.1% and incubated at 37 ℃. 10 µL aliquots were collected at 0, 1, and 3 hr and analyzed by MALDI-TOF MS after spotting with ⍺-cyano-4-hydroxycinnamic acid (Millipore Sigma).

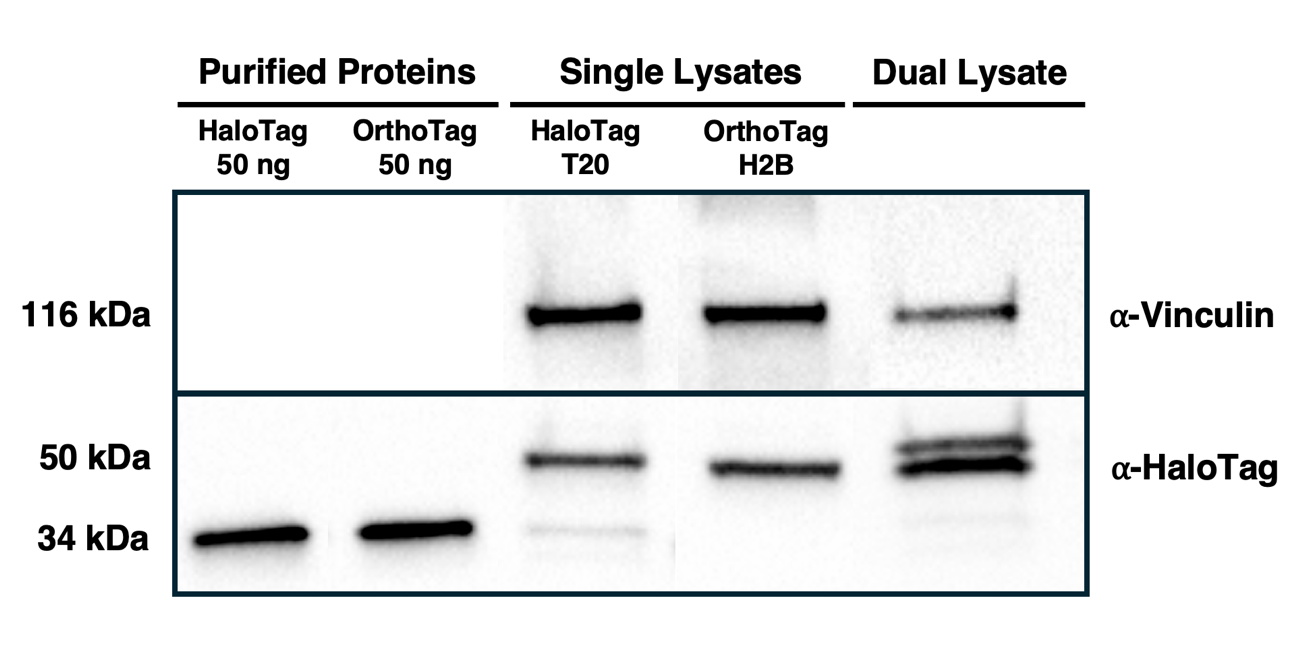

#### **Supplemental Figure 16. Expression levels of OrthoTag and HaloTag in mammalian cells.** The expression levels of OrthoTag-H2B and HaloTag-T20 in stable HEK293T cell lines expressing either each protein individually or both proteins together were evaluated by western blot as previously described.^8^ Briefly, 3 x 10^5^ cells were plated in a 6-well plate the day before an experiment. Cells were harvested by washing in PBS twice and incubating directly with 300 µL of ice-cold RIPA buffer (ThermoFisher Scientific) with 1x cOmplete™️ EDTA-free protease inhibitor cocktail (Millipore Sigma) and Pierce™️ Phosphatase Inhibitor Mini Tablets (ThermoFisher Scientific) before scraping and sonicating for 20 min in a bath sonicator. Cell debris was sheared through a 27G needle 3 times before pelleting at 4 ℃ for 3 min at 21,000 xg. Total protein concentration was measured by Pierce™️ 660 nm protein assay (ThermoFisher Scientific). 12 µg of protein was loaded into each lane of a 4-20% Mini-PROTEAN®️ TGX™️ Precast Protein Gels (BioRad) and electrophoresed before transferring to a 0.45 µm PVDF transfer membrane (ThermoFisher Scientific). Membranes were blocked with dry milk powder in TBST buffer (1x, pH 7.6, 0.1% tween-20) (ThermoFisher Scientific) before labeling with 1x ⍺-HaloTag mouse monoclonal primary antibody (Promega) or ⍺-vinculin mouse monoclonal primary antibody (Millipore Sigma) and goat ⍺-mouse-HRP polyclonal secondary antibody (Invitrogen), developing with SuperSignal™️ West Pico PLUS Chemiluminescent Substrate (ThermoFisher Scientific), and measuring luminescence using a Bio-Rad ChemiDoc™️ XRS+. The primary ⍺-HaloTag antibody was first validated using a known concentration of purified 6xHis OrthoTag and HaloTag to ensure that it labeled each protein equivalently.

### Confocal Fluorescence Microscopy and In-Gel Fluorescence

Colocalization analysis was performed using CellProfiler on a population of 20 adjacent cells (Fig. 4b,c).^9^ The BDY channel (corresponding to HaloTag7-TOMM20) was used to assign cell borders as objects. Colocalization was measured between the Hoechst, BDY, and TMR channels within cell objects with a threshold value set to 15% of the maximum intensity and the Pearson correlation coefficients were recorded (Fig. 4c).

#### **Supplemental Table 11. Confocal imaging acquisition parameters.**

| Instrument | Objective | λ_ex_ [nm] (%) | λ_em_ [nm] | Size [pxl] | Scan Speed [Hz] | Optical Zoom |
| --- | --- | --- | --- | --- | --- | --- |
| Leica DMi8 (SP8) | 40x water, 63x water | 405 (1)  488 (1)  552 (1) | 410-495  495-561  561-733 | 1024 | 600 | 1-2x |

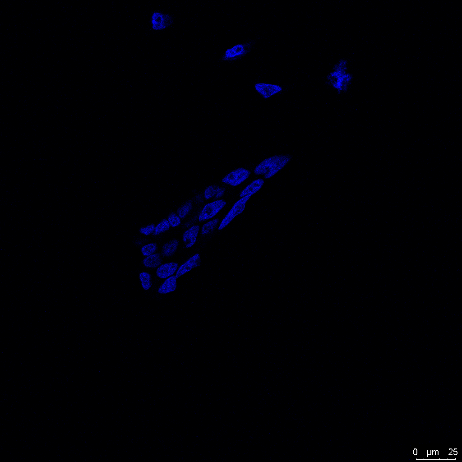

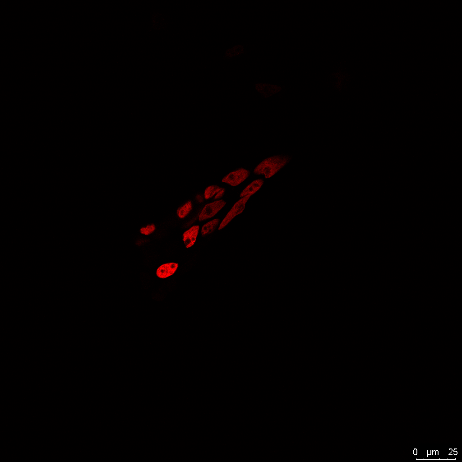

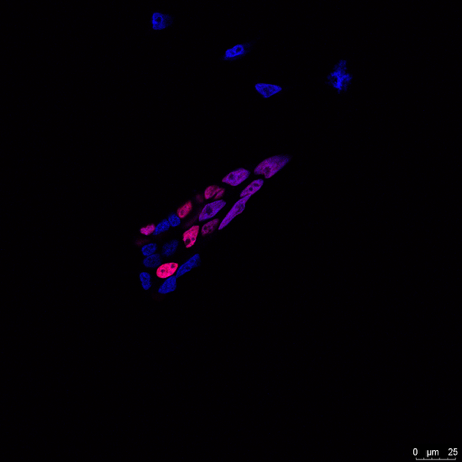

#### **Supplemental Figure 17. Microscopy images showing nuclear localization of OrthoTag-H2B fusion.** HEK293 cells expressing OrthoTag-H2B were labeled with 30 nM bCA-TMR for 1 h before washing, fixing, and staining with Hoechst 33342. The Hoechst channel is shown on the left, the TMR channel in shown in the middle, and the merged image is shown on the right.

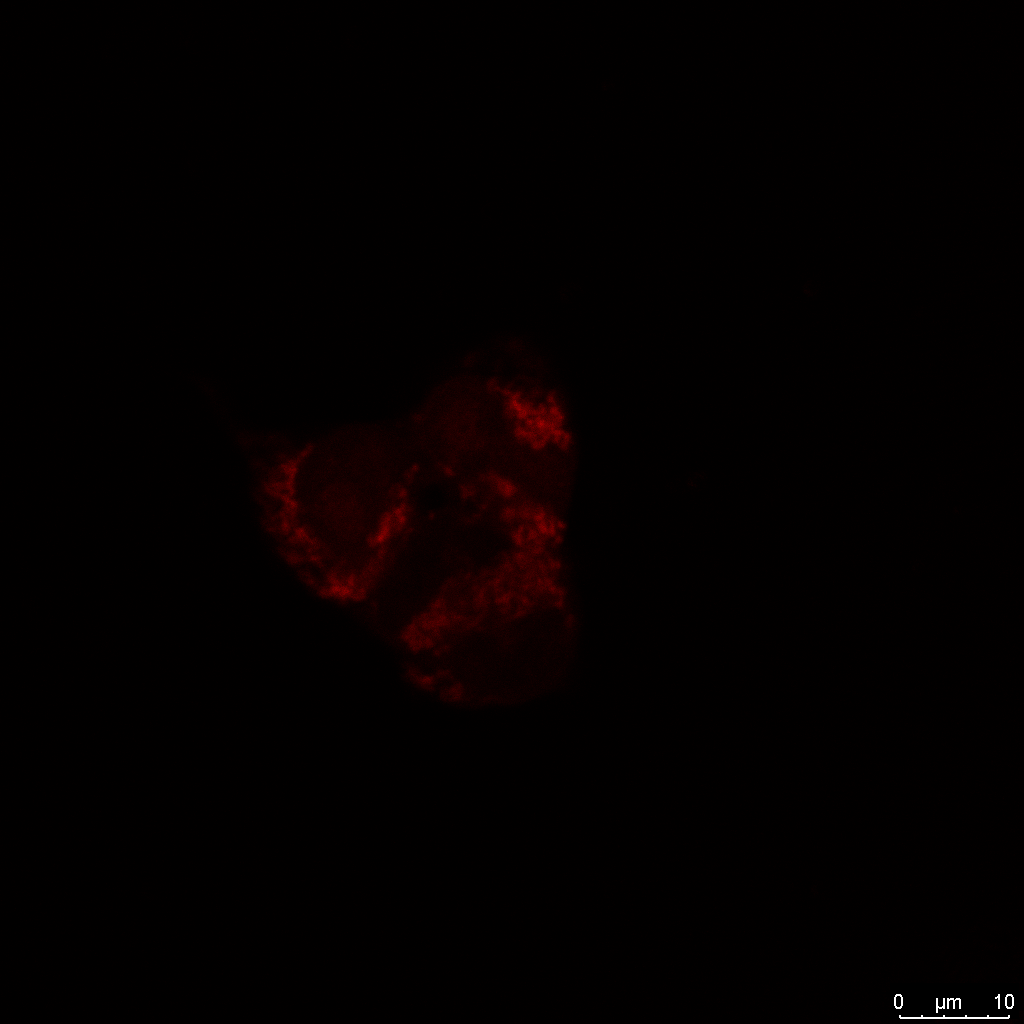

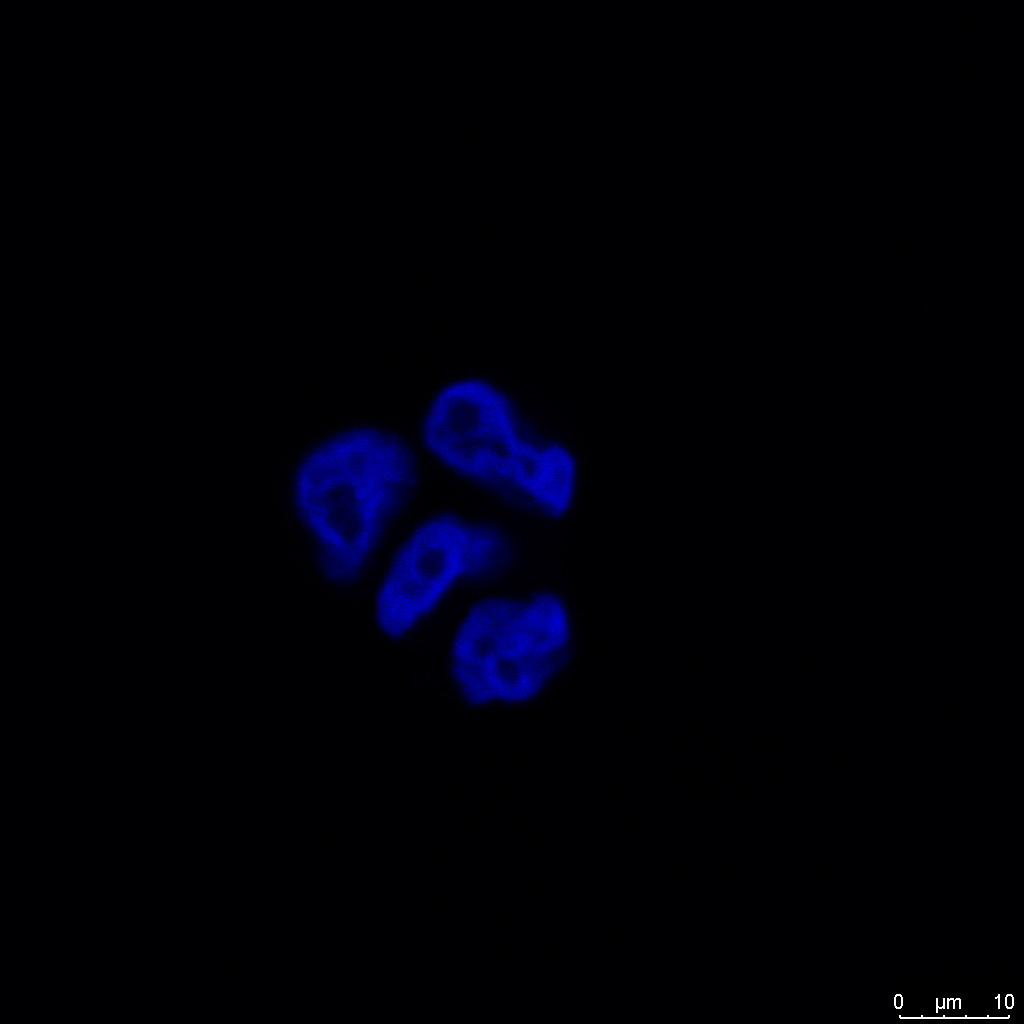

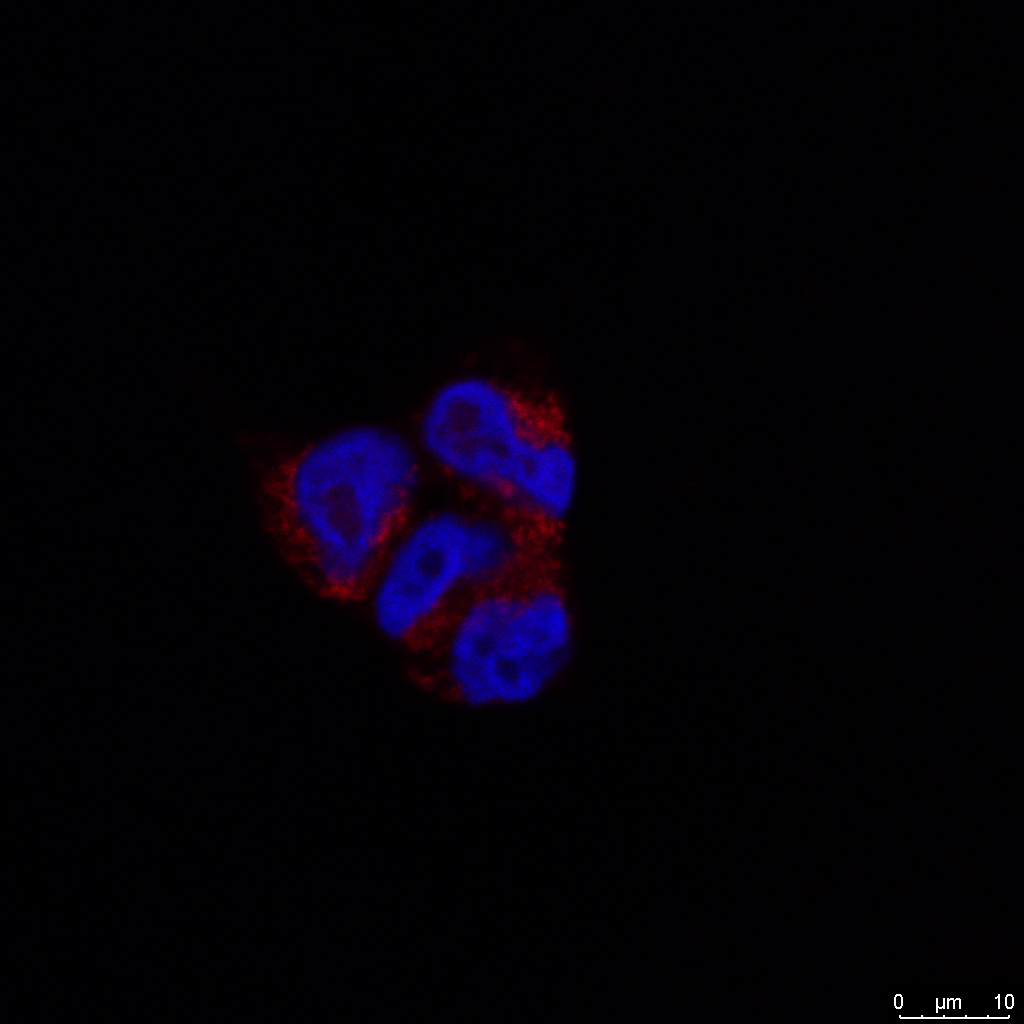

#### **Supplemental Figure 18. Mitochondrial localization.** HEK293T cells were stained with Hoechst and MitoTracker™️ DeepRed FM to visualize mitochondria following manufacturer protocols. The MitoTracker™️ DeepRed FM channel is shown on the left, the Hoechst channel is shown in the middle, and the merge of the two channels is shown on the right.

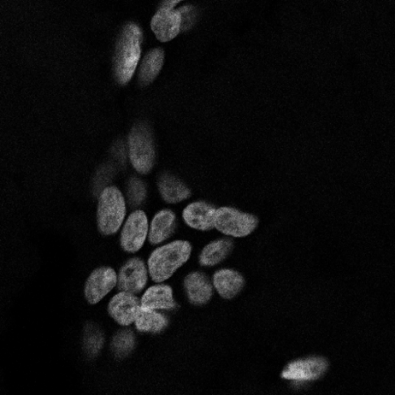

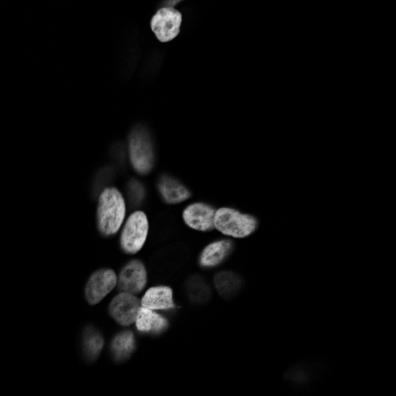

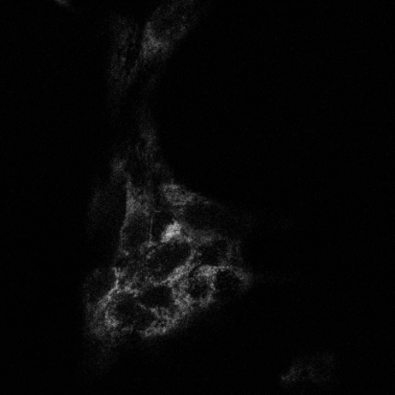

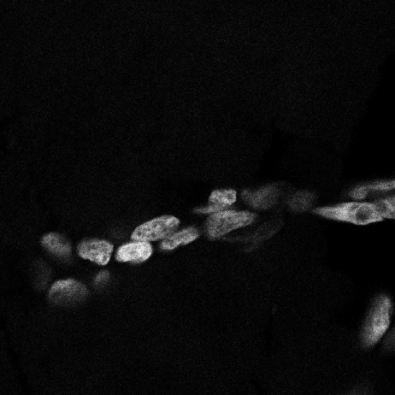

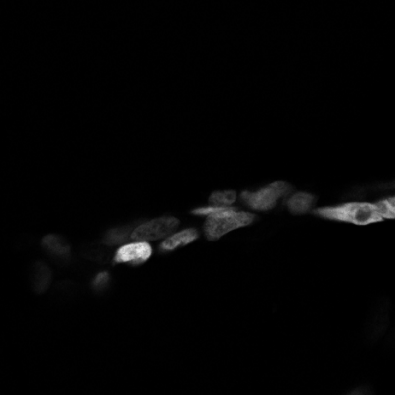

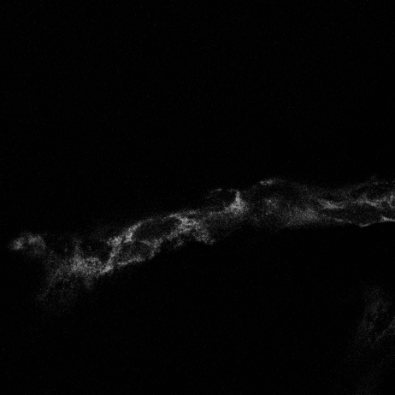

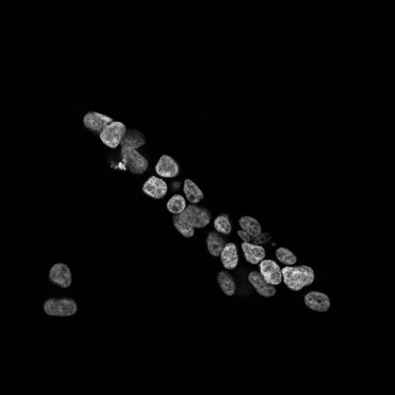

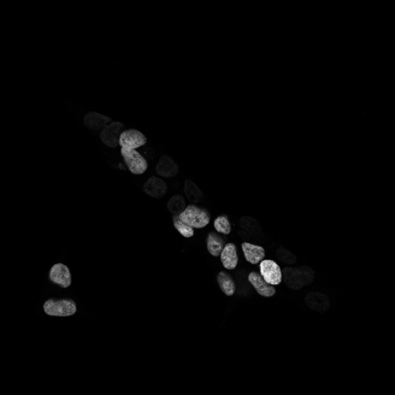

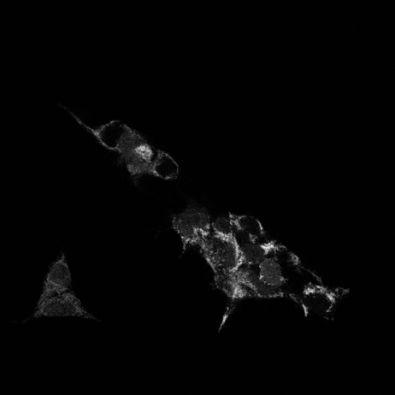

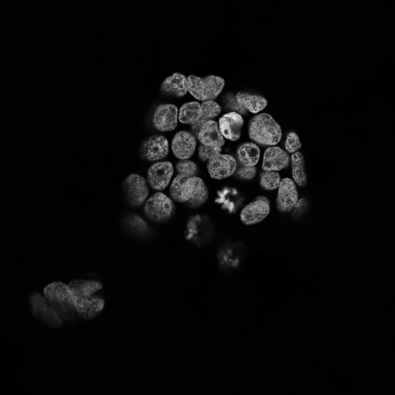

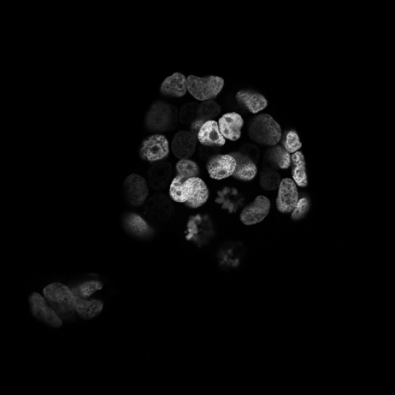

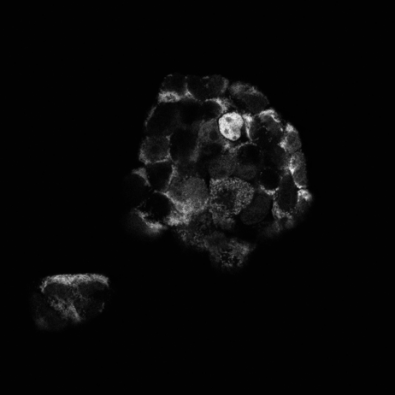

#### **Supplemental Figure 19. Representative raw microscopy images from multiplexed labelling experiment.** The top three images correspond to the colored panels in Fig. 4. The other images are from separate groups of cells under the same treatment and imaging conditions. HEK293T cells expressing both HaloTag-T20 and OrthoTag-H2B were labeled with 100 nM CA-BDY and 100 nM bCA-TMR for 1 h before washing, fixing, and staining with Hoechst 33342. The Hoechst channel is shown on the left, the TMR channel in shown in the middle, and the BDY channel is shown on the right.

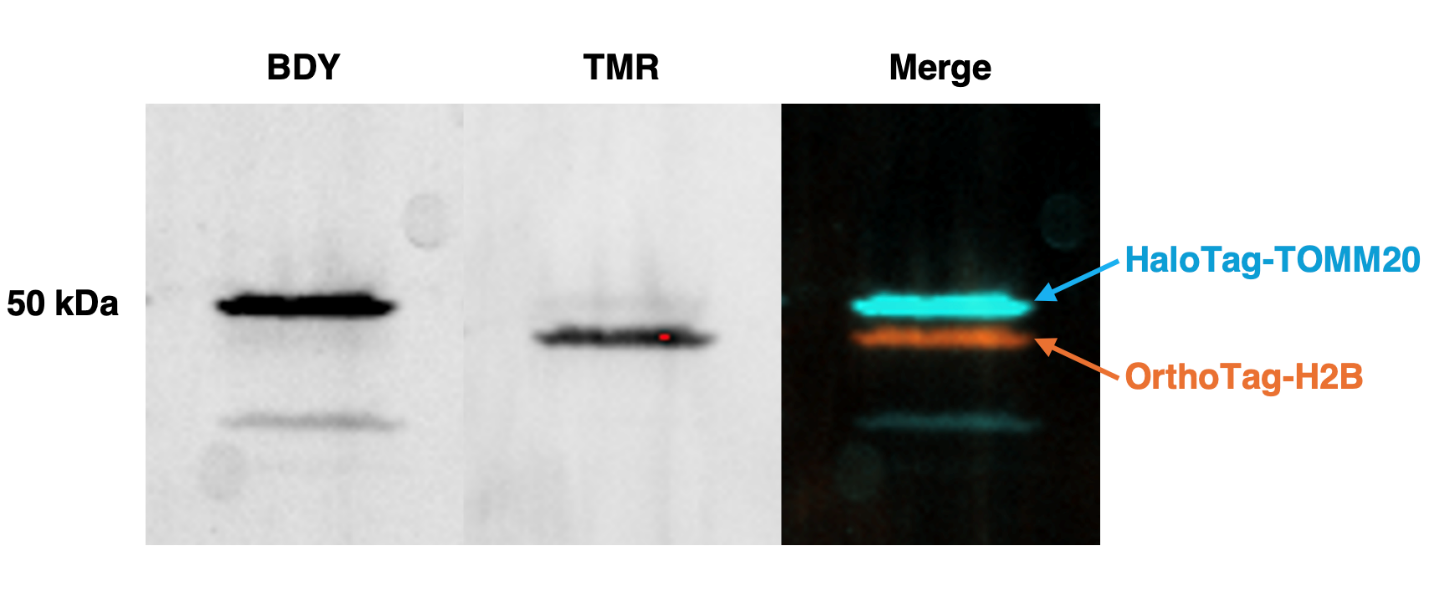

#### **Supplemental Figure 20. In-gel fluorescence confirms target engagement in cells.** Two days prior to the experiment, 3 x 10^5^ HEK293T cells stably expressing both OrthoTag-H2B and HaloTag-T20 were plated in a 6-well plate. These cells were then treated with 200 nM bCA-TMR and 50 nM CA-BDY in OPTI-MEM. bCA-TMR was added 30 min prior to CA-BDY. 30 min after CA-BDY was added, treatment was aspirated, and cells were washed twice with OPTI-MEM for 30 min at 37 ℃ and 5% CO_2_ in a humidified incubator. Before harvesting, cells were washed twice with cold PBS, and then total lysates were harvested, and 18 µg of total protein were electrophoresed as described in the western blotting protocol. Gels were imaged on an Invitrogen™️ iBright™️ 1500FL gel imager using the FITC and TRITC channels.

### General Methods and Instrumentation

All plasmids were amplified in 5⍺ competent *E. coli* (NEB) and purified by QiaPrep® Spin Miniprep Kit (Qiagen) or PureLink™️ HiPure Plasmid Maxiprep Kit (ThermoFisher Scientific). Plasmids were stored as pure stocks in nuclease free water at -20 °C or 20% glycerol stocks of 5⍺ *E. coli* at -80 °C.

All antibodies were purchased from ThermoFisher Scientific unless otherwise stated.

Spectral characterization was performed using a Bruker 400 MHz NMR and Thermo Finnigan LTQ MS. Masses of degradation and covalent labeling products were detected using a Bruker Autoflex maX MALDI. Fluorescence polarization assays and absorbance measurements were performed using a Tecan Spark plate reader. Flow cytometry experiments were performed on either a Cytek® Guava® easyCyte™ or an Agilent Novocyte Quanteon cell analyzer. A Bio-Rad S3e™ cell sorter was used for fluorescence-activated cell sorting. Confocal microscopy was conducted on a Leica FLIM SP8 and images were processed using the FIJI package of ImageJ and CellProfiler. Gel images were gathered on a Bio-Rad ChemiDoc™️ XRS+ or Invitrogen™️ iBright™️ 1500FL. Crystal structures were analyzed in PyMOL and using the POVME3 binding pocket analysis software.

#### **Supplemental Table 12.** **Primer list.**

| Primer Name | Description | Sequence (5’-3’) |
| --- | --- | --- |
| P1 | HaloTag fwd | ATGGCTGAAATTGGTACAGGTTTTCCATTTG |
| P2 | HaloTag rev | AGAAATTTCTAAAGTTGACAACCATCTAGCAATTTC |
| P3 | HaloTag pCTcon2 Gibson fwd | GGAGGCGGTAGCGGAGGCGGAGGGTCGGCTAGCTGCGGTGGCGGCGGTATGGCTGAAATTGGTACAGGTTTTCCATTTG |
| P4 | HaloTag pCTcon2 Gibson rev | GTCCTCTTCAGAAATAAGCTTTTGTTCGGATCCGCCCCCAGAAATTTCTAAAG TTGACAACCATCTAGCAATTTC |
| P5 | HaloTag pET Gibson fwd | AGAAGGAGATATACCATGCATCACCACCATCATCACATGGCTGAAATTGGTAC AGGTTTTC |
| P6 | HaloTag pET Gibson rev | AGCGGTGGCAGCAGCCTAGGTTAATTAAGAAATTTCTAAAGTTGACAACCATC |

#### **Supplemental Table 13.** **Standard PCR conditions.**

| Step | Temp (°C) | Time (s) | Cycles |
| --- | --- | --- | --- |
| Initial Denaturation | 95 | 60 | 1 |
| Denaturation | 95 | 30 | 35 |
| Annealing | *depends on primer* | 15 |  |
| Extension | 68 | 60 / kilobase |  |
| Final Extension | 68 | 300 | 1 |
| Hold | 4 | ∞ |  |

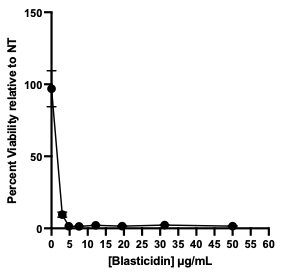

#### **Supplemental Figure 21. Blasticidin S kill curve for HEK293T cells.** Cell viability was measured using the MTT Cell Proliferation Assay (ATCC) following manufacturer protocols after ten days of culturing in blasticidin-supplemented media. 5000 cells were seeded in standard media a 96-well plate at day -1 and swapped into selective media on day 0. Selective media was changed every two days until day ten when major loss in viability was noted, and the MTT assay was performed to determine the lowest concentration of blasticidin that could be used for selections (5 μg/mL). Data points represent the average of three technical replicates with error bars representing the standard error of the mean.
